# A Highly Thermostable and Novel GH5 Endoglucanase from *Bacillus* sp. Strain BS with Enhanced Biomass Saccharification Potential in Seawater

**DOI:** 10.64898/2025.12.21.695781

**Authors:** Debjyoti Ghosh, Aditi Konar, Yi Qin Gao, Manas Mondal, Sushant K. Sinha, Supratim Datta, S. Venkata Mohan

## Abstract

The efficient enzymatic conversion of lignocellulosic biomass to glucose is crucial for biofuel production, with enzyme costs being a significant barrier to commercialization. Multifunctional enzymes that reduce enzyme components can improve efficiency. In this study, we amplified and sequenced a gene from *Bacillus* sp. strain BS, which showed 89.8% identity with bacterial endoglucanase Q6YK34. This putative novel endoglucanase, *Bs*EG2, is a GH5-like endoglucanase, with optimal activity at pH 6.0 and 55 □, and efficiently hydrolyzes amorphous and crystalline cellulose, including pre-treated and untreated sugarcane bagasse. Kinetics on carboxymethyl cellulose (CMC) indicate high substrate binding efficiency (apparent *V*_max_ of 148.9 µM min^−1^, a *K*_m_ of 24.9 mg mL^−1^, and a *k*_cat_ of 517 s^−1^). *Bs*EG2 hydrolyzes long cellulosic chains, oligosaccharides, and cellobiose to produce glucose, demonstrating multifunctionality. It exhibits remarkable thermal stability, retaining over 90% activity after 15 days at 55 °C. *Bs*EG2 also retains activity in high salts and ionic liquids, and is not inhibited by cellobiose, even at 200 mM, unlike other endoglucanases and cellobiohydrolases. Furthermore, *Bs*EG2 functions in seawater, reducing freshwater footprint in biofuel production. *Bs*EG2 is a promising candidate for a cost-effective enzyme formulation for glucose production from lignocellulosic biomass, supporting downstream glucose-fed metabolic pathways.

## Introduction

Lignocellulosic biomass is an abundant feedstock for biofuels, biomaterials, and chemicals; however, efficient cellulose-to-glucose conversion requires a synergistic action of a cellulase cocktail ^[1]^. Cellulose depolymerization typically involves three enzyme types: endoglucanases (EC 3.2.1.4), which randomly cleave β-1,4 glycosidic bonds to produce shorter oligosaccharides and expose new chain ends; cellobiohydrolases (EC 3.2.1.91), which act on the newly exposed ends to release cellobiose; and β-glucosidases, which hydrolyze cellobiose to glucose. Producing such cocktails is complex and costly, accounting for nearly 40 % of total bioethanol production costs ^[2]^. Therefore, the development of robust, cost-effective cellulases with high specific activity, tolerance to hydrolysis products, metal ions, salts, and ionic liquids, and long half-lives is critical.

An attractive alternative is multifunctional, processive cellulases that combine activities in a single enzyme. For example, bifunctional cellulases may combine the activities of endoglucanases and exoglucanases, while trifunctional cellulases may also include β-glucosidase activity. Several bifunctional cellulases have been identified, such as CelB from *Caldocellum saccharolyticum*, CelA from *Anaerocellum thermophilum*, and *Ct*Cel5E, a bifunctional cellulase/xylanase from *Clostridium thermocellum*^[3]^. However, to date, no naturally occurring trifunctional cellulase that yields glucose as the primary product has been reported. Processivity, the enzyme’s ability to continuously thread a cellulose chain through its active site to perform successive cleavages, enhances hydrolytic efficiency by generating oligosaccharides that are amenable to further conversion into glucose^4^.

Bacterial cellulases, especially those from *Bacillus* species, are attractive because these organisms tolerate diverse environmental conditions ^[4]^. *Bacillus subtilis* is particularly noteworthy for producing a variety of industrially relevant enzymes, including protease, amidase, and cellulase, which makes it a promising candidate for pharmaceutical applications ^[5]^. Certain strains of *B. subtilis*, such as KB-1111 and KB-1122, have been investigated as biocontrol agents and are known to secrete endo-1,4-β-glucanase ^[6]^.

Here, we report a novel GH5 family, endoglucanase-like gene from a newly isolated *Bacillus* sp. strain BS, encoding an enzyme designated *Bs*EG2 + CBM. We cloned two constructs into pET21b+. *Bs*EG2, containing the signal peptide and catalytic domain, and another including the signal peptide, catalytic domain, and a short carbohydrate-binding module (*Bs*EG2+CBM). Both constructs were heterologously expressed in *Escherichia coli* and purified for biochemical characterization. *Bs*EG2 exhibits multifunctional and processive behavior that challenges its classification as a conventional endoglucanase. Molecular dynamics simulations comparing the binding of cellooligosaccharide G6 to *Bs*EG2 with and without CBM reveal differences in substrate orientation and interaction energetics, suggesting the CBM influences substrate positioning at the catalytic site.

## Result and Discussion

### Identification of a novel endoglucanase-like gene

PCR and Sanger sequencing of *Bacillus sp*. strain BS yielded a full-length 1263 bp gene encoding the protein designated *Bs*EG2+CBM. The gene encodes a protein that contains a signal peptide, a catalytic domain, and a carbohydrate-binding module (CBM). Pairwise comparison to the endoglucanase Q6YK34 showed 88.7% nucleotide identity and 89.8% amino-acid identity, indicating a putative novel endoglucanase-like gene hereafter referred to as BsEG2.

### Cloning, expression, and purification of BsEG2

The catalytic-domain-only (BsEG2) and catalytic-domain-plus-CBM (BsEG2+CBM) coding sequences were cloned into pET-21b+ under a T7 promoter with an N-terminal His6 tag, and verified by Sanger sequencing (primers in Table S1). Both constructs were overexpressed in *E. coli* BL21(DE3) and purified from soluble extracts by HisTrap affinity chromatography. SDS–PAGE (12%) showed each purified protein as a single predominant band (*Bs*EG2, Figure S1a; *Bs*EG2+CBM, Figure S1b).

Domain and family assignments were performed using InterPro^[7]^ and PANTHER^[8]^. A conserved motif (VIYEIANEPN) maps to Glycoside Hydrolase family 5 (PTHR34142), InterPro annotations identify endoglucanase features and a CBM3a domain (Table S2), and multiple-sequence alignment with GH5 members revealed conserved catalytic and structural sequence stretches (Figures S2–S3).

### Optimum temperature, pH, and specific activity of *Bs*EG2

The pH and temperature dependence of *Bs*EG2+CBM and *Bs*EG2 was evaluated on the soluble substrate carboxymethyl cellulose (CMC). *Bs*EG2+CBM exhibited the highest activity at 60 °C within the pH range of 5.0 to 6.0 (Figures S4 and S5). *Bs*EG2, on the other hand, displayed an optimal temperature (T_opt_) of 55 °C, retaining over 90% activity between 50-65 °C (Figure 1a). At its T_opt_, *Bs*EG2 retained more than 80% of its specific activity within the pH range of 5.0 to 7.5, with peak activity observed at pH 6.0 – 6.5 in McIlvaine buffer (Figure 1b). These optimal conditions are consistent with previously reported endoglucanases from *Bacillus* sp^[9]^.

**Figure 1.**
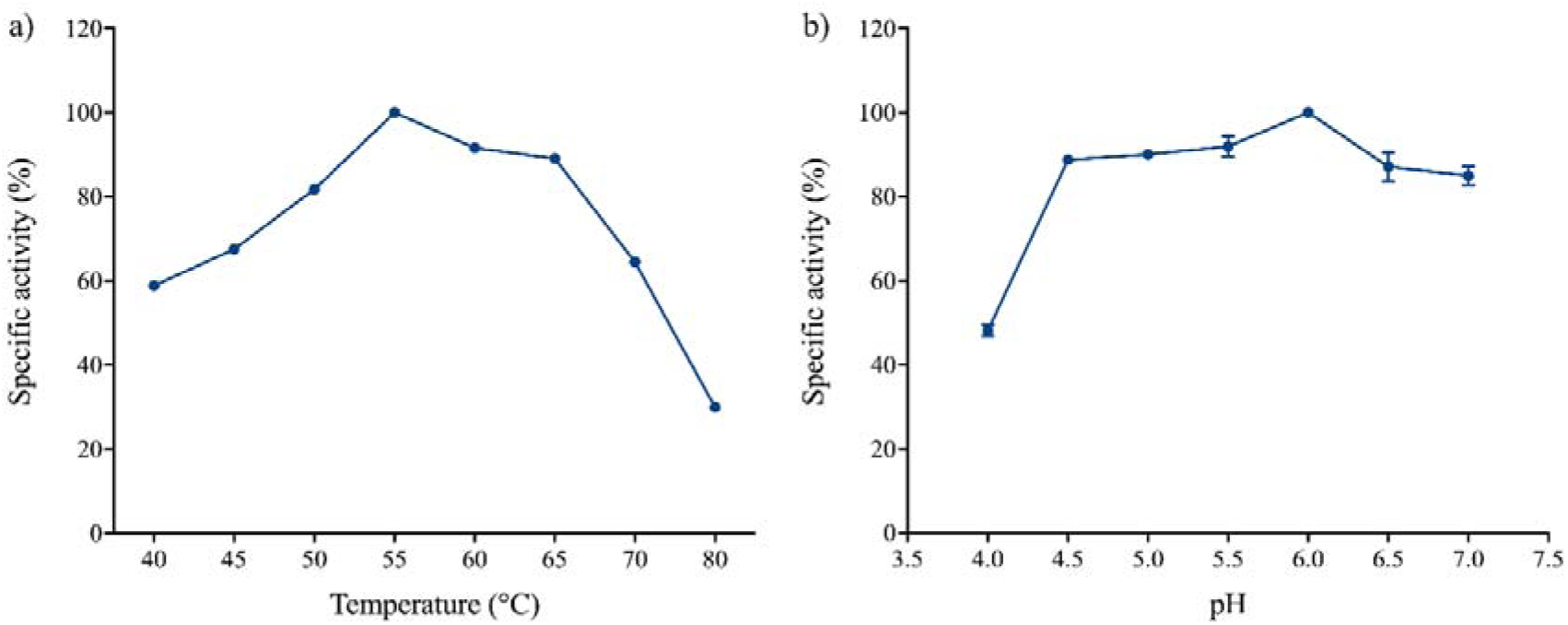
Determination of temperature and pH optima of *Bs*EG2: a) The effect of temperature on *Bs*EG2 activity was measured between 40 to 80 °C using 1 % CMC as the substrate in McIlvaine buffer, pH 6.0. Specific activity was measured using the DNS assay and reported as % specific activity. b) The effect of pH on *Bs*EG2 activity was determined by measuring specific activity in McIlvaine buffer across a pH range of 4 to 7 at the T_opt_ (55 °C) on 1 % CMC. 100 % specific activity denotes the BsEG2 specific activity (340.57 ± 10.61 μmoles min^−1^ mg^−1^) recorded at the optimum temperature and pH of *Bs*EG2. Error bars represent standard deviations from independent reactions.

Under its optimal conditions, pH_opt_ 6.0 and T_opt_ of 55 °C, *Bs*EG2 exhibited a specific activity of 340.6 ± 10.6 U mg-1 (where 1 U = 1 µmol of reducing sugars generated per minute of the reaction per mg of enzyme used in the enzymatic reaction). *Bs*EG2+CBM showed a specific activity of 327 ± 24.8 U mg^−1^ under its respective optimal conditions, indicating that the short CBM did not significantly alter activity on this soluble substrate. To further investigate the CBM’s role, enzymatic activity was tested on both untreated and pre-treated sugarcane bagasse. No significant difference was observed after CBM removal (Figure 2), in contrast to previous studies that highlight the importance of CBMs in binding insoluble substrates^[10]^. While these results suggest that the CBM does not enhance substrate catalysis in *Bs*EG2, further investigations are required to assess its potential role in thermostability or other functional properties. Enzymes may exhibit high optimal temperatures but shorter half-lives, or vice-versa, as observed in examples such as I7DLX2 from *Caldicellulosiruptor* sp. F32, P50388 from *Saccharolobus shibatae*, P22498 from *Saccharolobus solfataricus*, and D9PZ08 from *Acidilobus saccharovorans*^[11]^.. In this report, however, most experiments were carried out only with the catalytic domain (*Bs*EG2).

**Figure 2.**
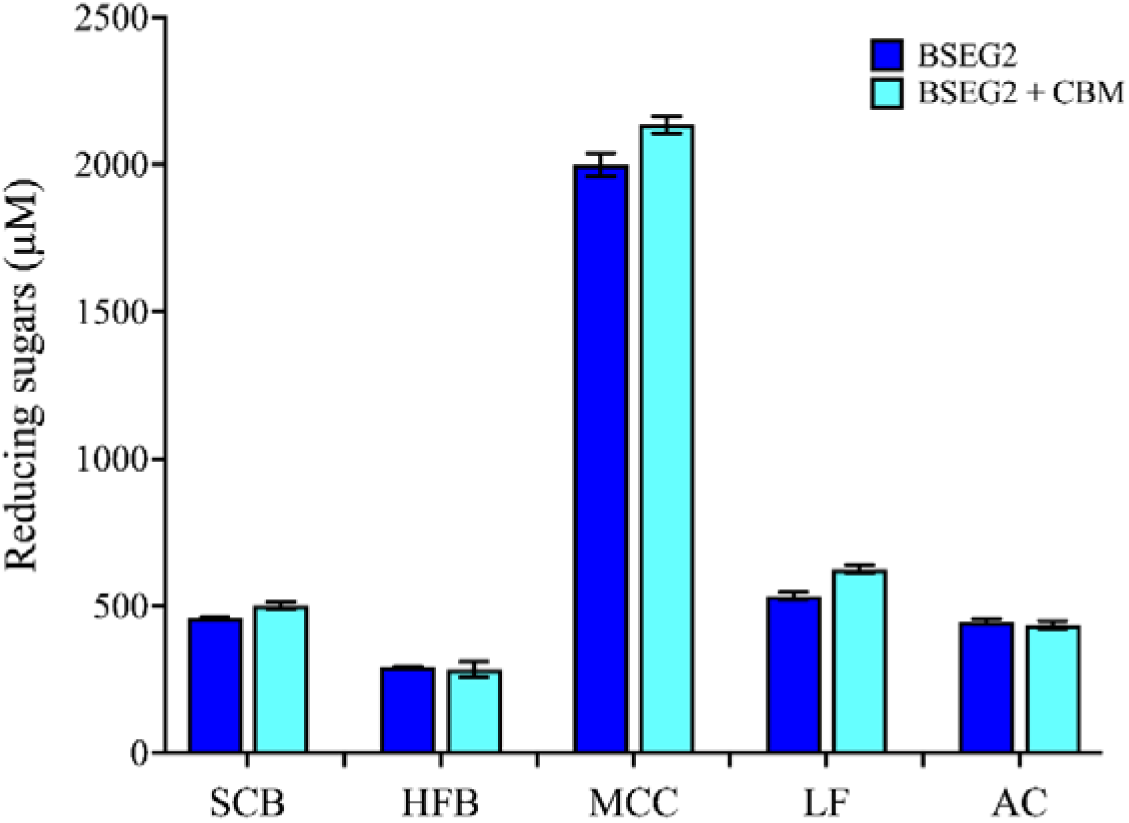
Hydrolytic activity of *Bs*EG2 and *Bs*EG2 + CBM on various biomass substrates, including SCB, sugarcane bagasse; HFB, hemicellulose-free biomass; MCC, microcrystalline cellulose; LF, lignin-free biomass; AC, α-cellulose; in the absence of carbohydrate-binding module. The hydrolysis reactions were performed for 2 hours using 5 µg enzyme under optimal temperature and pH. The resulting hydrolysis products were quantified using the DNS assay.

### Substrate specificity of *Bs*EG2

The kinetic parameters of *Bs*EG2 on CMC were determined under optimal assay conditions (pH 6.0, 55 °C) in McIlvaine buffer. Data were analyzed using the non-linear regression fit in GraphPad PRISM (version 8.0). Complete substrate saturation could not be achieved within the usable concentration limit of 24 mg mL^−1^, requiring the extrapolation of the curve. Similar cases of unsaturated Michaelis-Menten kinetics have also been reported for other enzymes, such as RsEG_m_ and EGL1^[12]^. On CMC, *Bs*EG2 has an apparent *V*_max_ of 148.9 µM min^−1^, an apparent *K*_m_ of 24.9 mg mL^−1,^ and a *k*_cat_ of 517 s^−1^ (Figure S6).

*Bs*EG2 substrate specificity was assessed using both natural and commercially available substrates (Table S3). The enzyme displays higher activity on β-glucan than on CMC, a trend observed in many endoglucanases ^[13]^. This preference is likely due to β-glucan’s greater solubility in water and its mixed β-1,3 and β-1,4 glycosidic linkages^[14]^. *Bs*EG2 also hydrolyzes insoluble crystalline substrates, such as Avicel-PH 101, filter paper, lichenan, and natural biomass, albeit at lower rates likely due to reduced accessibility and higher crystallinity^[15]^. In addition, *Bs*EG2 hydrolyzes chromogenic substrates, including azo-CMC, *p*NPClb, and *p*NPLac (Table S3). The enzyme can also act on natural cellulosic substrates, such as both pre-treated and untreated sugarcane bagasse. Sugarcane bagasse, a fibrous residue after sugarcane stalks are crushed to extract juice, is an abundant biomass in many parts of the world, including India. Among various substrates, *Bs*EG2 exhibits the highest activity on microcrystalline cellulose (MCC), followed by lignin-free bagasse (LF). The *Bs*EG2 activity on real biomass remains similar with or without CBM, as *Bs*EG2 and *Bs*EG2+CBM show comparable activity under identical reaction conditions. (Figure 2).

The effect of *Bs*EG2 on filter paper and sugarcane bagasse was further examined using scanning electron microscopy (Figure S7). After 12 hours at 55 °C, the SEM image shows that the untreated filter paper (control) has a smooth, uniform, and dense mesh of cellulose fibers. Upon *Bs*EG2 treatment, this network became undulated, more open and non-uniform, and thinner, with interstitial fibres becoming more susceptible to enzymatic digestion. These results suggest that *Bs*EG2 can act on diverse cellulosic substrates and may have the potential to convert agricultural residues, such as sugarcane bagasse, into value-added products.

### Molecular docking and MD simulation to understand the role of CBM

The AlphaFold-predicted model structure of *Bs*EG2 reveals a structured catalytic domain and a short CBM (Figure S8(a)). The catalytic domain primarily consists of α-helices and β-sheets. Using molecular modelling, docking, and all-atom MD simulations, we further investigate the binding site of cellohexaose (G6) and its interactions with the *Bs*EG2 in the presence and absence of CBM. Cellohexaose (G6) was used as a structural representative equivalent of long cellulosic chains for the *in silico* docking studies. The docked structures of *Bs*EG2 with G6 exhibited the highest docking free energy scores of –7.97 kcal/mol with CBM (Figure 3(a)) and –6.74 kcal/mol without CBM (Figure 3(b)). By analysing the distances between receptor-ligand interface residue pairs that are below 5.0 Å, we identified the key *Bs*EG2 residues involved in forming bonded and non-bonded contacts with G6 (Table S4). Notably, the binding site residues remain mostly similar in both scenarios, though the orientation of G6 differs at the binding site pocket depending on the presence of CBM.

**Figure 3:**
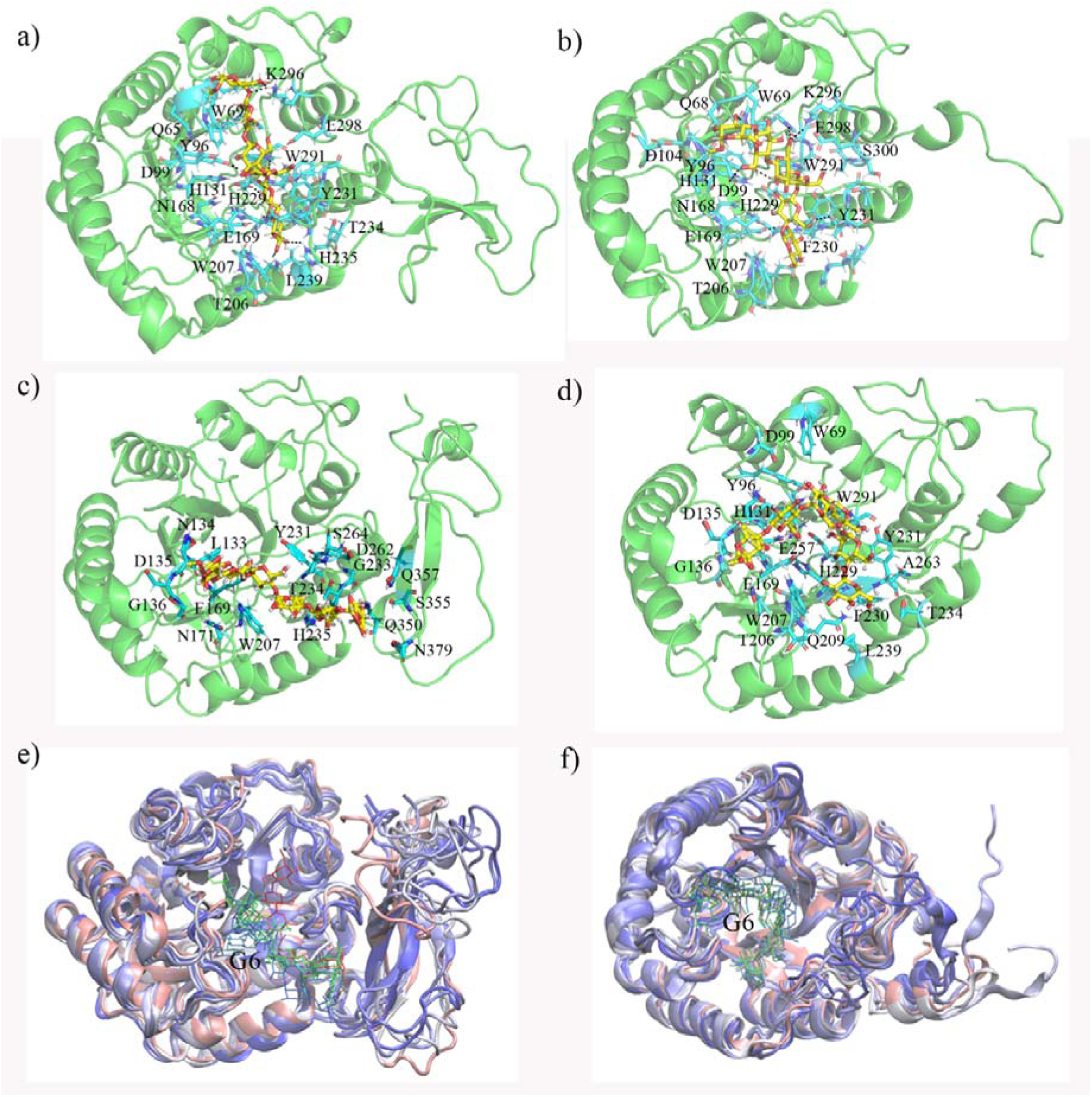
(a)-(b) Initial and (c)-(d) simulated average G6 docked structures of *Bs*EG2 with and without CBM. The representative G6 binding site residues and G6 are shown as stick representations and marked with cyan and yellow colors, respectively. (e)-(f) The overlay structures of G6 bound *Bs*EG2 with and without CBM along the equilibrated MD trajectories, where the conformation of protein and ligand in different simulation times are marked with RWB and RGB color scale.

Furthermore, we evaluated the equilibrium dynamics of *Bs*EG2, the stability of the G6 at the binding site pocket, and the energetics of G6 binding through microsecond-long MD simulations. Over the equilibrated trajectory, G6 stabilizes the binding site pocket of *Bs*EG2 in different orientations, relative to the initial docked conformation (Figure S8(b)-(c)). The convergence of the root-mean-square displacement (RMSD) of the G6-bound *Bs*EG2 structures with respect to their initial energy-minimized structures over the simulation time (Figure S9(a)) suggests that G6 can form stable complexes with BsEG2 both in the presence and absence of CBM. Notably, G6 is positioned deeper within the catalytic domain’s binding pocket of *Bs*EG2 when CBM is absent. The distance between the mass center of G6 and the catalytic domain of *Bs*EG2 (d_G6-catalytic_) increases in the presence of CBM, as illustrated in Figure S9(b). Figure 3(c)-(d) shows the simulated average docked structures of G6 along with representative binding site residues of *Bs*EG2 with and without CBM. The structural overlay of G6-bound *Bs*EG2 along the equilibrated MD trajectories (Figure 3(e)-(f)) reveals distinct preferred binding conformations of G6 within the binding pocket, along with the flexible movements of G6 when CBM is present (Figure 3(e)). Conversely, G6 shows restricted movements with lower root mean square fluctuations (RMSF) of the sugar moieties in the absence of CBM compared to the *Bs*EG2 with CBM (Figure S9(c)). On average, five favorable polar hydrogen-bonded contacts can form between G6 and *Bs*EG2 (Figure S9(d)). Key residues forming stable polar contacts with the oxygen atoms of G6 in the binding pocket of *Bs*EG2 without CBM include (Table S4) Y96, D99, N134, W207, Q209, Y231, and E257. In contrast, in the presence of CBM, residues L133, E169, G233, T234, and D262 from the catalytic domain establish favorable polar contacts with G6. Most residues, within the binding pocket of *Bs*EG2 that participate in the G6 binding (Table S4), are highly conserved (Figure S10).

Both electrostatic (E_elec_) and van der Waals (E_vdw_) interactions favor G6 binding with *Bs*EG2, where the E_vdw_ is more favorable for *Bs*EG2 without CBM (Table S4). We further evaluated the free energy of binding G6 to *Bs*EG2 and the contributions of individual binding site residues using the MMGBSA approach ^[16]^. The negative total binding free energy (Δ*G*), as indicated in Table S4, points to a favorable interaction between G6 and *Bs*EG2, regardless of the presence of CBM. However, the estimated Δ*G* values do not accurately represent the actual binding free energy due to the omission of entropic factors related to the release of solvent molecules. The residues contributing significantly to the favorable ΔG for G6 binding configurations with and without CBM are listed in Table S4. Notably, residues such as L133, W207, Q209, H229, F230, F231, T234, E257 and W291 are highly conserved (Figure S10). Overall, our *in-silico* results characterize the binding site and suggest that the binding of G6 to *Bs*EG2 and *Bs*EG2+CBM is favorable. Additionally, it also reveals differential conformational flexibility of substrate G6 within the binding site pocket of *Bs*EG2 in the presence and absence of CBM, suggesting further roles for CBM in processive polysaccharide translocation. In the substrate-bound state, the flexible movement of G6 within the catalytic pocket of *Bs*EG2 is correlated with the flexible motions of CBM, which might be important for processivity. Further studies are needed to fully understand the finer implications of the role of the truncated CBM.

### Stability of *Bs*EG2

Ionic liquids (ILs) are excellent non-aqueous solvents for lignocellulosic biomass pretreatment^[17]^. However, residual ILs can inhibit cellulase activity, necessitating extensive washing, which increases water consumption and costs^[18]^. IL-tolerant enzymes are therefore desirable for compatibility with IL-pretreated biomass and for probing determinants of enzyme stability. *Bs*EG2 was evaluated for its tolerance to imidazolium-based ILs, specifically 1-ethyl-3-methylimidazolium [C_2_mim] variants (Figure 4a). It maintained full activity in up to 20% [C_2_mim][Cl], 15% [C_2_mim][OAc], and 10% [C_2_mim][Lac], though higher concentrations of [C_2_mim][OAc] and [C_2_mim][Lac] resulted in 20% and 40% activity loss, respectively. This suggests that anion identity plays a critical role in enzyme stability. ILs typically inhibit cellulase activity via denaturation, aggregation, competitive inhibition, or modification in enzyme dynamics^[18b, 19]^. The surface charge of enzymes has also been proposed to play a role in maintaining the integrity and specific activity in ionic liquids^[20]^. Notably, *Bs*EG2 contains 24% (92 out of 383) charged residues, among the highest reported for endoglucanases, which may contribute to IL tolerance.

**Figure 4.**
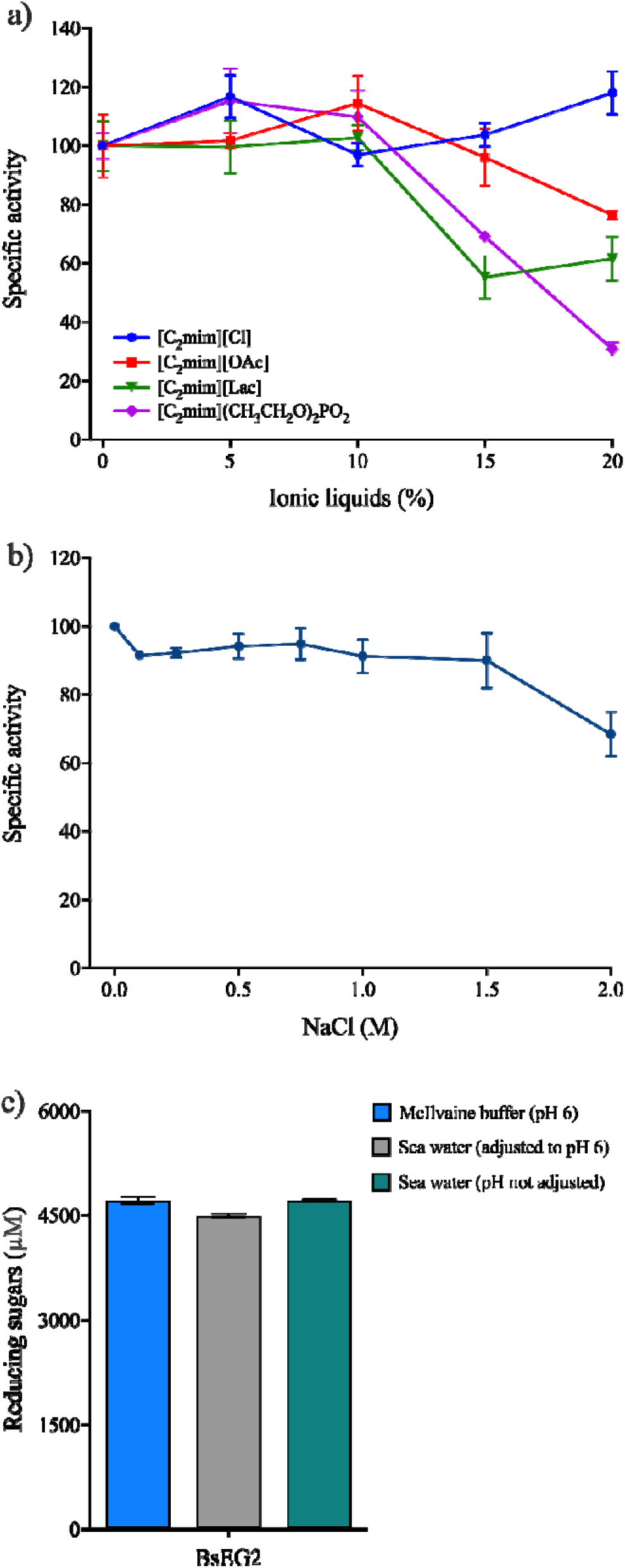
a) Effect of the [C_2_mim] cation-based ionic liquid (IL) on B*s*EG2 activity. Specific activity was measured in the presence of 5 to 20% IL using 1% β-glucan as the substrate in McIlvaine buffer, pH 6.0. Residual specific activity was determined using the DNS assay and expressed as a percentage of the enzyme’s activity in the absence of ionic liquid (set at 100%). **b)** Salt tolerance of *Bs*EG2. The enzyme was incubated in McIlvaine buffer at pH_opt_ and 55°C with 0.2 to 2 M NaCl, and then assayed under standard conditions. The specific activity of *Bs*EG2 in the absence of NaCl was set at 100% specific activity (340.57 ± 10.61 U/mg). Error bars represent the standard deviation from three independent experiments. **c)** Comparison of reducing sugar concentrations produced by *Bs*EG2 and *Bs*EG2+CBM in McIlvaine buffer (at pH 6), seawater (pH adjusted to 6), and seawater (pH unadjusted) at 60°C. The reducing sugars generated in each reaction were quantified using the DNS assay.

*Bs*EG2 also shows high salt tolerance, retaining over 90% activity up to 1.5 M NaCl and around 70% activity at 2 M NaCl, though the salt tolerance is not as high as in some previously reported EGs ^[21]^. (Figure 4b). Since pre-treated biomass often contains residual salts from acid or base neutralization, such tolerance can reduce the need for extensive washing, lowering overall process costs. Salt-tolerant cellulases may also allow the substitution of freshwater with non-potable sources, such as seawater, in bioprocessing operations^[22]^. *Bs*EG2 functions effectively in seawater at its natural pH (7.8-8.1) and pH 6.0, indicating its suitability for marine or seaweed biomass degradation without pretreatment (Figure 4c).

Metal ion tolerance further supports *Bs*EG2’s industrial potential (Figure S11). Divalent (Ca^2+^, Cu^2+^, Mg^2+^, Ni^2+^, Zn^2+^) and trivalent Fe^3+^ cations slightly enhanced its activity by up to 25%, whereas monovalent cations (K^+^, Na^+^) had no effect. This resilience to metal contaminants commonly present in feedstocks or process water increases BsEG2’s applicability *Bs*EG2 thermal stability is outstanding. After 12 days at 55 °C, *B*sEG2 retained 59 ± 2 % activity, corresponding to an approximate half-life of 15 days (Figure 5), and retained 97 ± 3 % activity after two months at 4 °C. The *Bs*EG2 melting temperature (T_m_) is 65.5 °C, classifying it as a thermophilic enzyme^59^. Enzymes with high T_m_ often favor prolonged high-temperature saccharification and reduced contamination risk, as they resist denaturants better than mesophilic enzymes^[23]^. In summary, *Bs*EG2 combines IL and salt tolerance, long half-life, tolerance to metal ions, and thermostability, making it an excellent candidate for biorefinery applications.

**Figure 5.**
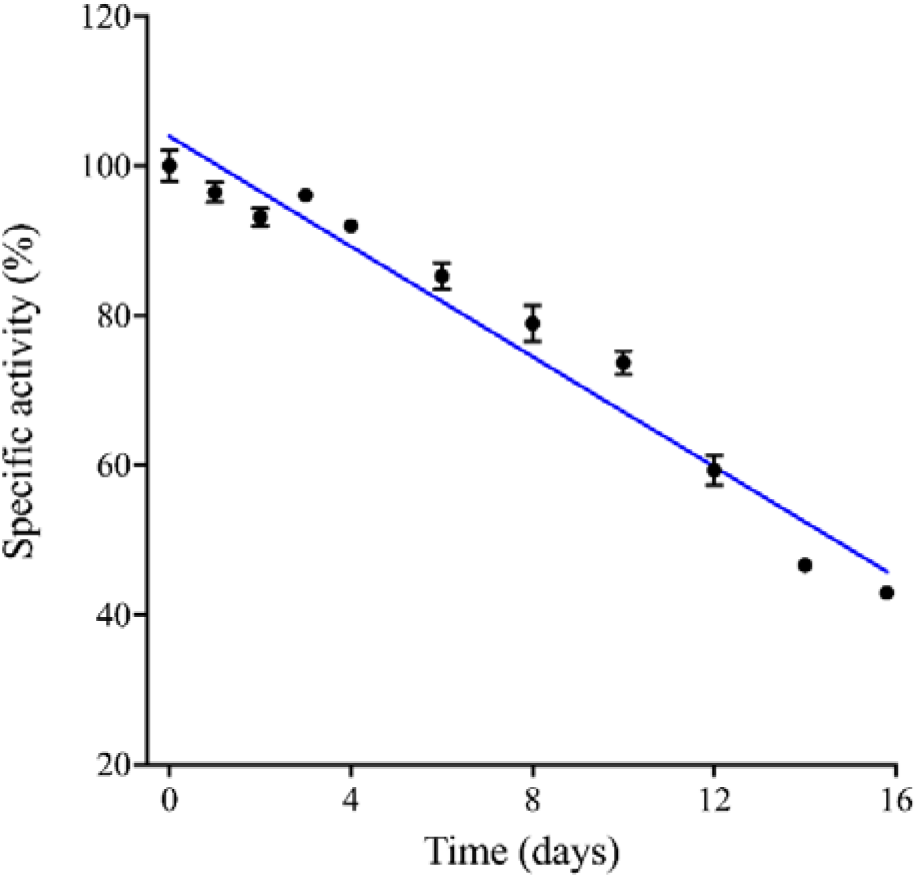
Thermostability of *Bs*EG2. The enzyme was incubated in buffer at pH_opt_ (pH 6.0) and 55 □C for up to 16 days. At designated time points, aliquots were taken and assayed under standard kinetic assay conditions. The specific activity of *Bs*EG2 at 0 h of incubation (340.57 ± 10.61 U/mg) was set as 100 %. Error bars represent the standard deviation from independent reactions.

### Processivity in *Bs*EG2

Hydrolysis by *Bs*EG2 yielded 2.6-fold more reducing sugars in the soluble fraction than in the insoluble fraction (Figure 6), indicating a processive mode of action. This behavior contrasts with typical endoglucanases, which often generate more reducing ends in the insoluble particles than in the soluble fraction^[24]^. Processivity is crucial in cellulose hydrolysis and is quantified by the ratio of soluble to insoluble reducing sugars. A higher processivity ratio indicates increased production of reducing sugars in the soluble fraction. While most processive endoglucanases belong to the GH9 family^[25]^, several GH5 members are also processive (e.g.cel5H from *Saccharophagus degradans*, p4818Cel5_2A from porcine gut microbiota, Cel5A from *Gloeophyllum trabeum*, CHU_2103 from *Cytophaga hutchinsonii*, and EG5C-1 from *Bacillus subtilis* BS-5 ^[9a, 13c, 26]^. Table S5 summarizes the processivity across the GH5 family.

**Figure 6.**
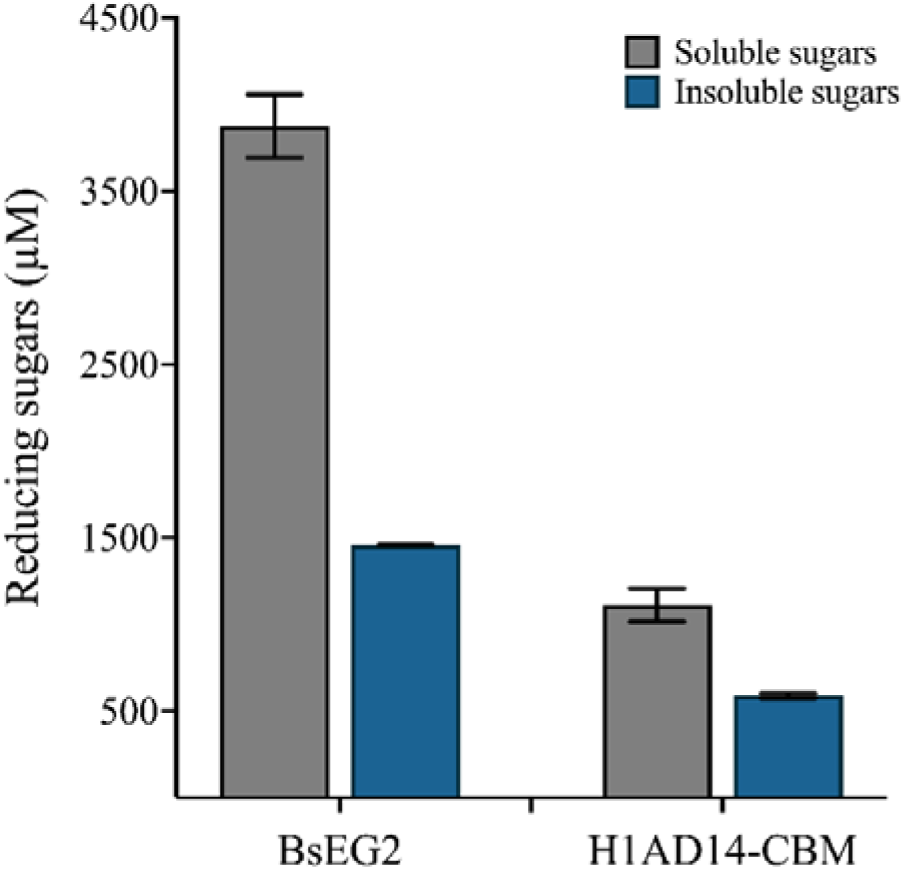
The ratio of soluble to insoluble reducing sugars generated from 30 mg of filter paper by 15 μg of enzyme in 3 hours at the enzyme’s optimal temperature and pH was determined using the DNS assay. The soluble-to-insoluble sugar ratio of *Bs*EG2 was compared to that of another processive endoglucanase, H1AD14, which loses most of its activity in the absence of its CBM. Reactions were performed in triplicate. Error bars represent the standard deviation from independent reactions.

Figure 6 contrasts the soluble and insoluble fractions of reducing sugars produced by *Bs*EG2 with those generated by the GH9 endoglucanase H1AD14-CBM (only catalytic domain). Unlike *Bs*EG2, the removal of the CBM in H1AD14 lowered its ratio of soluble to insoluble reducing sugars, whereas the presence of CBM3 enhanced its processivity^[26a, 27]^. The processivity in GH5 endoglucanases does not always depend on the presence of CBMs. For example, deleting the N-terminal CBM1 in the endoglucanase from *Volvariella volvacea* affected its processivity, whereas the deletion of CBM6 in GH5 endoglucanases (*Sd*Cel5G, *Sd*Cel5H, and *Sd*Cel5J) from *Saccharophagus degradans* had no impact^[26c, 28]^. Additionally, endoglucanases from *Gloeophyllum trabeum* and *Cytophaga hutchinsonii* have been reported to be processive even without CBMs^[26b, 26c]^. CBM3, which is natively present in many GH9 endoglucanases, has been shown to enhance processivity^[29]^.

### Trifunctionality of *Bs*EG2

The enzymatic activity of *Bs*EG2 was further evaluated using a range of substrates, including oligosaccharides. As reported earlier (*vide infra*), *Bs*EG2 hydrolyzed chromogenic substrates like *p*NPClb and *p*NPLac, which are structural analogues of trisaccharide, generating cellobiose, lactose, and *p*NP as hydrolysis products. To further analyze its substrate specificity, we assessed *Bs*EG2 hydrolysis of cello-oligosaccharides after 10-minute and 30-minute incubations by HPLC (Figure 7a, Figure S12). The hydrolysis products of cellopentaose (G5) resulted in the formation of cellotetraose (G4), cellotriose (G3), and cellobiose (G2). After 10 minutes, residual cellotetraose was present, but it became negligible after 30 minutes, coinciding with an increased accumulation of cellobiose. Similarly, the cellotetraose hydrolysis after 10 min yielded cellotriose, cellobiose, and some unreacted cellotetraose. After 30 minutes, cellotetraose was nearly depleted, and the accumulation of cellobiose had doubled. A comparable trend was observed in the hydrolysis of cellotriose (Figure S12). The higher accumulation of cellobiose compared to cellotriose at both time points suggests that *Bs*EG2 preferentially produces cellobiose.

**Figure 7.**
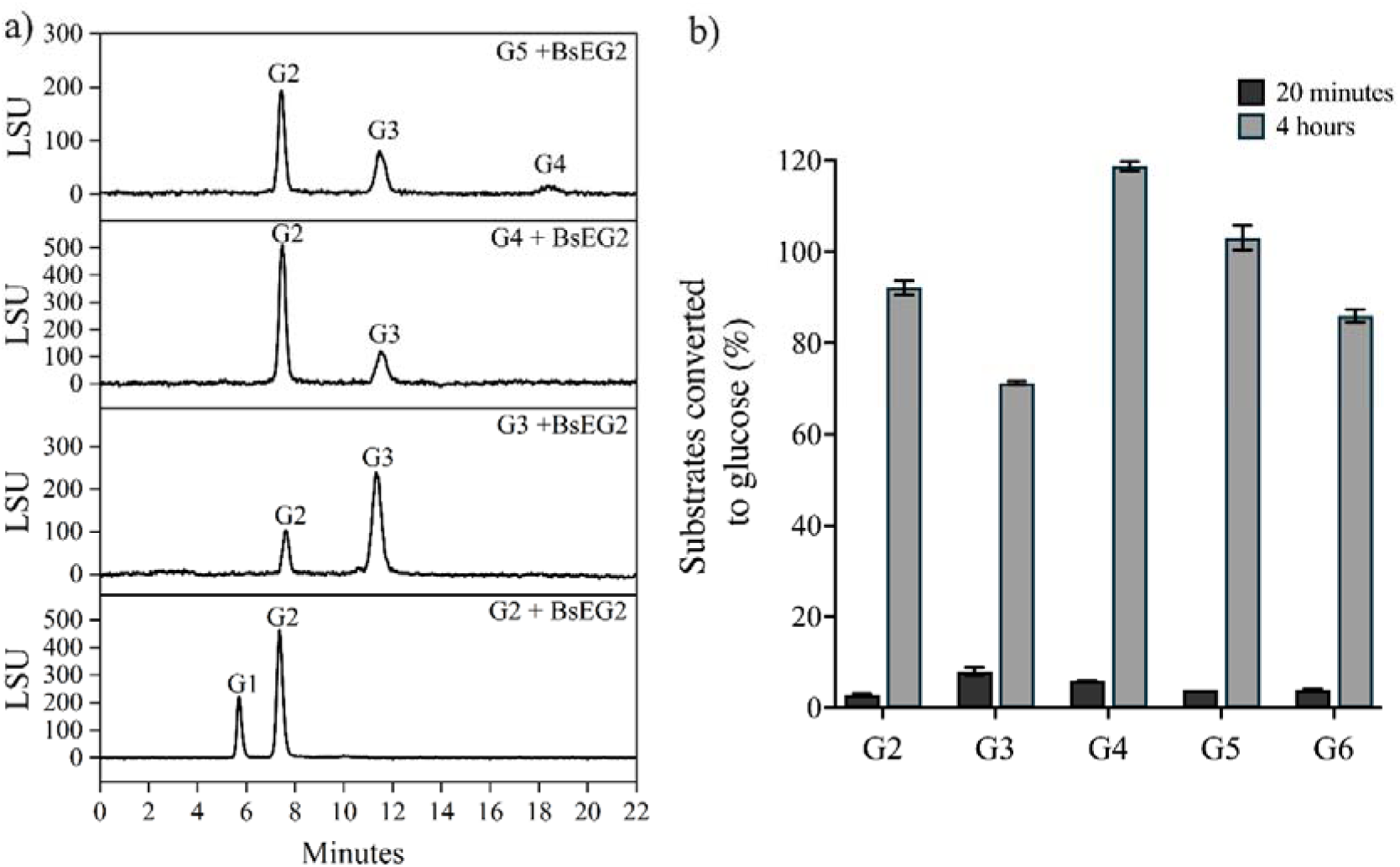
(a) HPLC analysis of *Bs*EG2 reaction products after 10 minutes of incubation with oligosaccharides. The assay was performed as described in the Methods section. The HPLC chromatogram displays the hydrolysis products of cellopentaose (G5), cellotetraose (G4), cellotriose (G3), and cellobiose (G2). **(b)**. Quantification of glucose production from various oligosaccharides (G2, cellobiose; G3, cellotriose; G4, cellotetraose; G5, cellopentaose; G6, cellohexaose) over with time. Within 4 hours, nearly 100% of the oligosaccharides were converted to glucose by 10 μM *Bs*EG2 at the optimal temperature and pH, as determined by the GOD-POD assay. Error bars represent the standard deviation from independent reactions.

However, when the reaction time was extended to four hours, all the oligosaccharide substrates, including cellobiose, cellotriose, cellotetraose, cellopentaose, and cellohexaose, were completely converted to glucose (Figure 7b). The conversion was quantified by GOD POD assay, and TLC analysis confirmed that glucose was the sole product, as indicated by a single spot in each reaction sample (Figure S13). Furthermore, when *Bs*EG2 was incubated with cellobiose for 20 minutes, HPLC analysis confirmed its hydrolysis to glucose (Figures 7a and S14). Notably, while some enzymes exhibit higher processivity than *Bs*EG2, they primarily generate cellobiose and cellotriose as end products, with only trace amounts of glucose being produced. Previous studies have detected minimal glucose formation by GH5 endoglucanases, such as in *AHK119-bMs* and *AHK119-E5* from *Thermobifida alba* AHK119, *Rf*GH5_4 from *Ruminococcus flavefaciens*, and *celM* from a thermal spring metagenome ^[30]^. In contrast, *Bs*EG2 exhibits a significantly higher glucose yield, suggesting a distinct hydrolytic mechanism that warrants further investigation.

Multifunctional cellulases offer a promising strategy for reducing enzyme costs in biomass saccharification by simplifying enzyme cocktail formulations and enhancing hydrolysis efficiency ^[31]^. *Bs*EG2, even though it is a multifunctional enzyme, also shares some sequence similarities with other endoglucanases within the GH5 family. While a sequence alignment of *Bs*EG2 with other GH5 endoglucanases revealed scattered stretches of conservation, the overall sequence similarity was limited, underscoring a distinct divergence and unique functional profile (Figures S1 and S2). Earlier attempts to develop single multidomain cellulases involved fusing two distinct cellulases via small linker peptides ^[32]^. However, such engineered multidomain fusions often resulted in misfolding and reduced expression or yield. In contrast, wild-type BsEG2 naturally combines endoglucanase, cellobiohydrolase, and β-glucosidase activities, enabling the direct conversion of cellulose to glucose (albeit at a slower rate than dedicated β-glucosidases) and offering potential for simplifying enzyme cocktails.

To understand why BsEG2 can produce significant amounts of glucose, we compared it to a putative endoglucanase from *Bacillus velezensis* (*B. velezensis*) (unpublished data), which has approximately 92% structural and sequence similarity with *Bs*EG2, but is unable to produce glucose. Analysis of the solvent accessible surface areas (SASA) and binding pocket volume of both enzymes using Discovery Studio revealed that their SASA values are nearly identical: 427.31 Å² for the endoglucanase from *B. velezensis* and 431.28 Å^2^ for *Bs*EG2. However, the volume of the substrate binding pocket is smaller in *Bs*EG2 (3,068.78 Å³ vs. 14,059.46 Å³ for *B*. *velezensis*). Tighter packing may constrain chain mobility in the cleft, favoring the longer residence of small substrates (e.g., cellobiose) and promoting further hydrolysis to glucose. Ongoing dynamic simulations and time-resolved studies of oligosaccharide binding and catalysis aim to clarify how pocket geometry and residue dynamics enable BsEG2’s pronounced glucose production.

### *Bs*EG2 has no inhibiting effects from reaction products

BsEG2 shows no inhibition by reaction products. During lignocellulosic biomass hydrolysis at high substrate concentrations, hydrolysis products such as oligosaccharides, cellobiose, and glucose can accumulate and inhibit enzyme activity, lowering saccharification efficiency^[33]^. Increasing enzyme loading to counteract this inhibition raises biofuel production costs. Among these hydrolysis products, cellobiose has been reported to cause stronger inhibition of cellulase activity than glucose^[34]^. To test *Bs*EG2’s tolerance, we measured its hydrolysis rate in the presence of added cellobiose and other hydrolysis products using Azo-CMC as substrate. BsEG2 retained full specific activity at cellobiose concentrations up to 200 mM (Figure S15). This product tolerance suggests that *Bs*EG2 may help overcome cellulase inhibition, thereby improving the efficiency and cost-effectiveness of biomass saccharification.

### Synergistic interactions of *Bs*EG2 with cellulase

Synergistic cooperation among cellulases is essential for efficient hydrolysis of long cellulose chains to glucose. We evaluated the synergy between *Bs*EG2 and the β-glucosidase B8CYA8 from *Halothermothrix orenii* on insoluble substrate Avicel (2% w/v), measuring glucose yield from combined enzyme action. The total enzyme load was kept constant to allow for a direct comparison of the impact of individual enzymes and the combined effects. The degree of synergy was calculated as the fold increase in glucose production relative to *Bs*EG2 alone.

*Bs*EG2 and B8CYA8 exhibit complementary activities. *Bs*EG2 primarily generates cellobiose, which B8CYA8 efficiently converts to glucose, enhancing overall cellulose hydrolysis (Figure 8). Glucose production from Avicel is reported as a percentage relative to *Bs*EG2 alone, which is considered 100%. The calculated degree of synergy between *Bs*EG2 and B8CYA8 is 1.6, indicating a cooperative effect on Avicel rather than competition for Avicel within the time periods of the assay, and the product of BsEG2 is utilized as substrate by B8CYA8.

**Figure 8.**
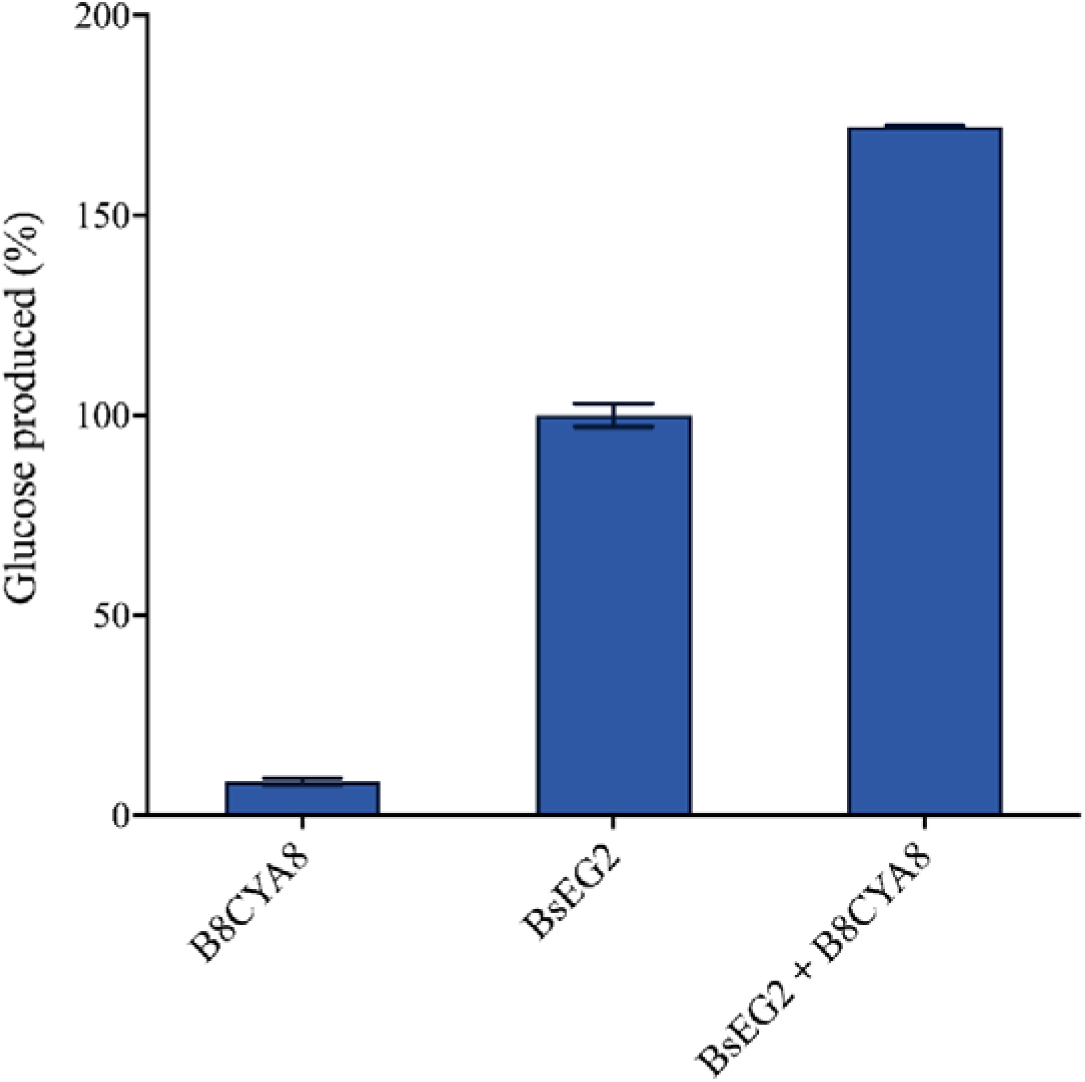
The synergistic effect of endoglucanase *Bs*EG2 with β-glucosidase (V169C/I246A/E173L B8CYA8) was evaluated on 2 % Avicel in McIlvaine buffer, pH 6.0, at 60 °C for 2 hours. Glucose concentration was quantified using the GOD-POD assay and expressed as a relative percentage, with the glucose yield from BsEG2 alone set at 100%.

An efficient cellulase cocktail design requires highly active enzymes and balanced functionalities. While BsEG2 can produce glucose on its own over longer time scales, combining it with BsEG2 yields faster glucose production via synergistic interaction.

## Conclusions

Despite originating from a mesophilic soil bacterium, *Bs*EG2, a GH5 endoglucanase, is active across a broad temperature and pH range, covering both mesophilic and thermophilic conditions. *Bs*EG2 is a highly efficient, processive endoglucanase from the GH5 family that efficiently hydrolyzes both amorphous and crystalline cellulose, as well as natural biomass such as sugarcane bagasse. Although classified as an endoglucanase, *Bs*EG2 can cleave long cellulose chains and even disaccharides, such as cellobiose, directly producing glucose. It has a long half-life and shows high tolerance for imidazolium-based ionic liquids and metal ions. *Bs*EG2 also retains activity in seawater (pH 5.0-7.0) and functions synergistically with glucose-tolerant β-glucosidases. It is not inhibited by reaction products. *In silico* analyses reveal that cellohexaose forms a stable complex within the catalytic pocket of BsEG2, stabilized by conserved polar and hydrophobic interactions, as well as flexible conformational dynamics that facilitate substrate binding and processive translocation. These combined properties make *Bs*EG2 a promising candidate for industrial biorefinery applications, justifying further study of its substrate specificity and mechanism.

## TOC figure

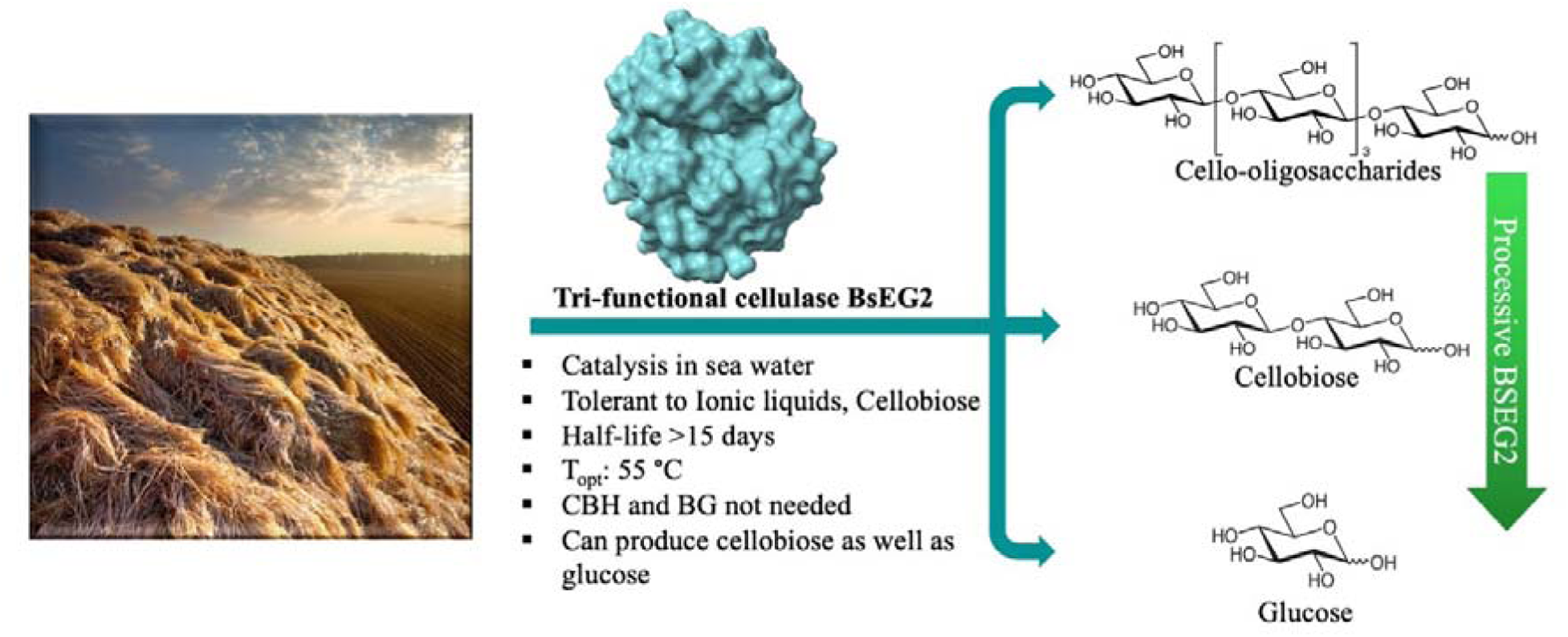

## References

[1] S. Datta, Current Metabolomics 2016, 4, 14–22.

[2] aD. Klein-Marcuschamer, P. Oleskowicz-Popiel, B. A. Simmons, H. W. Blanch, Biotechnol Bioeng 2012, 109, 1083–1087; bM. D. N. Ramos, T. S. Milessi, R. G. Candido, A. A. Mendes, A. Aguiar, Energy for Sustainable Development 2022, 68, 103–119.

[3] aS. J. Han, Y. J. Yoo, H. S. Kang, The Journal of biological chemistry 1995, 270, 26012–26019; bD. J. Saul, L. C. Williams, R. A. Grayling, L. W. Chamley, D. R. Love, P. L. Bergquist, Appl Environ Microbiol 1990, 56, 3117–3124; cV. Zverlov, S. Mahr, K. Riedel, K. Bronnenmeier, Microbiology (Reading, England) 1998, 144 (Pt 2), 457–465; dS. F. Yuan, T. H. Wu, H. L. Lee, H. Y. Hsieh, W. L. Lin, B. Yang, C. K. Chang, Q. Li, J. Gao, C. H. Huang, M. C. Ho, R. T. Guo, P. H. Liang, The Journal of biological chemistry 2015, 290, 5739–5748.

[4] R. Lynd Lee, J. Weimer Paul, H. van Zyl Willem, S. Pretorius Isak, Microbiology and Molecular Biology Reviews 2002, 66, 506–577.

[5] Y. Su, C. Liu, H. Fang, D. Zhang, Microbial Cell Factories 2020, 19, 173.

[6] C. X. Zhang, X. Zhao, F. Han, M. F. Yang, H. Chen, T. Chida, S. H. Shen, Journal of microbiology and biotechnology 2009, 19, 351–357.

[7] M. Blum, A. Andreeva, L. C. Florentino, S. R. Chuguransky, T. Grego, E. Hobbs, B. L. Pinto, A. Orr, T. Paysan-Lafosse, I. Ponamareva, G. A. Salazar, N. Bordin, P. Bork, A. Bridge, L. Colwell, J. Gough, D. H. Haft, I. Letunic, F. Llinares-López, A. Marchler-Bauer, L. Meng-Papaxanthos, H. Mi, D. A. Natale, C. A. Orengo, A. P. Pandurangan, D. Piovesan, C. Rivoire, C. J. A. Sigrist, N. Thanki, F. Thibaud-Nissen, P. D. Thomas, S. C. E. Tosatto, C. H. Wu, A. Bateman, Nucleic Acids Res 2025, 53, D444–d456.

[8] P. D. Thomas, D. Ebert, A. Muruganujan, T. Mushayahama, L. P. Albou, H. Mi, Protein science: a publication of the Protein Society 2022, 31, 8–22.

[9] aB. Wu, S. Zheng, M. M. Pedroso, L. W. Guddat, S. Chang, B. He, G. Schenk, Biotechnology for Biofuels 2018, 11, 20; bS. Regmi, Y. S. Choi, Y. K. Kim, M. M. Khan, S. H. Lee, S. S. Cho, Y.-Y. Jin, D. Y. Lee, J. C. Yoo, J.-W. Suh, Biotechnology and Bioprocess Engineering 2020, 25, 104–116; cW. Li, W.-W. Zhang, M.-M. Yang, Y.-L. Chen, Molecular Biotechnology 2008, 40, 195–201; dL. Lin, X. Meng, P. Liu, Y. Hong, G. Wu, X. Huang, C. Li, J. Dong, L. Xiao, Z. Liu, Applied Microbiology and Biotechnology 2009, 82, 671–679.

[10] A. B. Boraston, D. N. Bolam, H. J. Gilbert, G. J. Davies, The Biochemical journal 2004, 382, 769–781.

[11] aD.-D. Meng, Y. Ying, K.-D. Zhang, M. Lu, F.-L. Li, Molecular BioSystems 2015, 11, 3164–3173; bN. Y. Park, J. Cha, D. O. Kim, C. S. Park, Journal of microbiology and biotechnology 2007, 17, 454–460; cS. D’Auria, A. Morana, F. Febbraio, C. Vaccaro, M. De Rosa, R. Nucci, Protein Expression and Purification 1996, 7, 299–308; dV. M. Gumerov, A. L. Rakitin, A. V. Mardanov, N. V. Ravin, Archaea 2015, 2015, 978632.

[12] aP. Zhang, X. Yuan, Y. Du, J. J. Li, BMC biotechnology 2018, 18, 35; bW. Xiong, J. K. Yang, F. Y. Chen, Z. G. Han, Enzyme Microb Technol 2017, 97, 71–81.

[13] aS. Voget, H. L. Steele, W. R. Streit, Journal of Biotechnology 2006, 126, 26–36; bC.-J. Duan, L. Xian, G.-C. Zhao, Y. Feng, H. Pang, X.-L. Bai, J.-L. Tang, Q.-S. Ma, J.-X. Feng, Journal of Applied Microbiology 2009, 107, 245–256; cW. Wang, T. Archbold, J. S. Lam, M. S. Kimber, M. Z. Fan, Scientific Reports 2019, 9, 13630.

[14] M. J. Edney, B. A. Marchylo, A. W. MacGregor, Journal of the Institute of Brewing 1991, 97, 39–44.

[15] L. Zhu, J. P. O’Dwyer, V. S. Chang, C. B. Granda, M. T. Holtzapple, Bioresour Technol 2008, 99, 3817–3828.

[16] S. Genheden, U. Ryde, Expert opinion on drug discovery 2015, 10, 449–461.

[17] R. P. Swatloski, S. K. Spear, J. D. Holbrey, R. D. Rogers, Journal of the American Chemical Society 2002, 124, 4974–4975.

[18] aM. B. Turner, S. K. Spear, J. G. Huddleston, J. D. Holbrey, R. D. Rogers, Green Chemistry 2003, 5, 443; bA. Schindl, M. L. Hagen, S. Muzammal, H. A. D. Gunasekera, A. K. Croft, Frontiers in Chemistry 2019, 7.

[19] B. Manna, P. Chanda, S. Datta, A. Ghosh, The journal of physical chemistry. B 2023, 127, 8406–8416.

[20] aP. R. Burney, E. M. Nordwald, K. Hickman, J. L. Kaar, J. Pfaendtner, Proteins: Structure, Function, and Bioinformatics 2015, 83, 670–680; bL. B. Johnson, S. Park, L. P. Gintner, C. D. Snow, Journal of Molecular Catalysis B: Enzymatic 2016, 132, 84–90.

[21] S. Aich, R. K. Singh, P. Kundu, S. P. Pandey, S. Datta, Biotechnol Biofuels 2017, 10, 135.

[22] S. K. Sinha, M. Datta, S. Datta, Green Chemistry 2021, 23, 7299–7311.

[23] aS. Datta, B. Holmes, J. I. Park, Z. Chen, D. C. Dibble, M. Hadi, H. W. Blanch, B. A. Simmons, R. Sapra, Green Chemistry 2010, 12, 338–345; bS. Konar, S. K. Sinha, S. Datta, P. K. Ghorai, Journal of Molecular Liquids 2020, 306, 112879; cS. Aich, S. Datta, Applied Microbiology and Biotechnology 2020, 104, 3935–3945.

[24] V. V. Zverlov, N. Schantz, W. H. Schwarz, FEMS Microbiology Letters 2005, 249, 353–358.

[25] A. Konar, S. Aich, R. Katakojwala, S. Datta, S. V. Mohan, Appl Microbiol Biotechnol 2022, 106, 6059–6075.

[26] aB. J. Watson, H. Zhang, A. G. Longmire, Y. H. Moon, S. W. Hutcheson, Journal of Bacteriology 2009, 191, 5697–5705; bR. Cohen, M. R. Suzuki, K. E. Hammel, Applied and Environmental Microbiology 2005, 71, 2412–2417; cC. Zhang, Y. Wang, Z. Li, X. Zhou, W. Zhang, Y. Zhao, X. Lu, Applied Microbiology and Biotechnology 2014, 98, 6679–6687.

[27] A. I. Chiriac, E. M. Cadena, T. Vidal, A. L. Torres, P. Diaz, F. I. Javier Pastor, Applied Microbiology and Biotechnology 2010, 86, 1125–1134.

[28] F. Zheng, S. Ding, Applied and Environmental Microbiology 2013, 79, 989–996.

[29] A. Pakarinen, M. O. Haven, D. T. Djajadi, A. Várnai, T. Puranen, L. Viikari, Biotechnol Biofuels 2014, 7, 27.

[30] aT. Ohta, H. Horie, A. Matsu-Ura, F. Kawai, Journal of bioscience and bioengineering 2019, 127, 554–562; bP. V. Gavande, K. Kumar, J. Ahmed, A. Goyal, Int J Biol Macromol 2023, 224, 1395–1411; cN. Joshi, G. Kaushal, S. P. Singh, Biotechnol Bioeng 2021, 118, 1531–1544.

[31] E. Glasgow, K. Vander Meulen, N. Kuch, B. G. Fox, Current Opinion in Biotechnology 2021, 67, 141–148.

[32] aN. Adlakha, S. Sawant, A. Anil, A. Lali, S. S. Yazdani, Applied and Environmental Microbiology 2012, 78, 7447–7454; bS. D. Kurniasih, A. Alfi, D. Natalia, O. K. Radjasa, Z. Nurachman, Microbiological Research 2014, 169, 725–732.

[33] P. Andrić, A. S. Meyer, P. A. Jensen, K. Dam-Johansen, Biotechnology advances 2010, 28, 308–324.

[34] M. Holtzapple, M. Cognata, Y. Shu, C. Hendrickson, Biotechnology and Bioengineering 1990, 36, 275–287.

