## Supplementary Files for "A Highly Thermostable and Novel GH5 Endoglucanase from *Bacillus* sp. Strain BS with Enhanced Biomass Saccharification Potential in Seawater"

**Material and methods**

**Chemicals:** All chemicals used were of analytical grade or higher quality and were purchased from Merck (Darmstadt, Germany), Sigma-Aldrich (St. Louis, USA), and MP Biomedicals (Irvine, USA). Among the substrates, sodium carboxymethyl cellulose (CMC), Avicel PH-101, and other chromogenic substrates were purchased from Sigma-Aldrich (St. Louis, USA). β-Glucan from barley, oligosaccharides of glucose, and azo-carboxymethyl cellulose were obtained from Megazyme (Bray, Ireland). Ionic liquids and ionic salts were also from Sigma-Aldrich (St. Louis, USA). The media for cell culture growth were purchased from HiMedia Chemicals (Mumbai, India). The enzymes and reaction buffers were purchased from New England Biolabs (Ipswich, USA). Salt-free oligonucleotides were synthesized by GCC Biotech Pvt Ltd (Kolkata, India). 10 kDa cut-off Amicon Ultra-4 membranes (EMD Millipore, Billerica, USA) were used to pool the active fractions post-purification. Plasmid and gel extractions were performed using the commercial kits QIAquick Spin Miniprep Kit and QIAquick Gel Extraction Kit (QIAGEN, Hilden, Germany). The plasmid concentrations were measured using TECAN Infinite^®^ 200 Pro (TECAN Trading AG, Switzerland).‬‬

‬‬‬‬‬‬‬‬‬‬‬‬‬‬‬‬‬‬‬‬‬‬‬‬‬‬‬‬‬‬‬‬‬‬‬‬‬‬‬

**PCR amplification and identification of novel endoglucanase-like gene:** From the genomic DNA of *Bacillus* sp. strain BS, the forward and reverse primers (Table S1) were designed based on the protein, Q6YK34 (EMBL: AAN07019.1, *Bacillus subtilis* strain: ATTCAU195, Protein Id: AAN07019.1), targeting the endoglucanase, which amplified the gene corresponding to the proteins *Bs*EG2 and *Bs*EG2+CBM. The PCR amplicon was sequenced in both directions using the Sanger sequencing approach. The generated sequence data were subjected to BLASTX against the INSDS database and submitted to NCBI. The accession number of the generated sequence data is PX237215.

**Plasmids and strains:** The sequenced endoglucanase was cloned into a T7-bacterial expression plasmid pET21b+ (Agilent Technologies, Santa Clara, USA) using *Eco*RI-HF and *Xho*I restriction enzymes. *Escherichia coli* Top10F’ (Life Technologies, La Jolla, USA) was used as a cloning host. *E. coli* BL21(DE3) (ThermoFisher Scientific, Waltham, USA) was used for *Bs*EG2 expression.‬‬‬‬‬‬‬‬‬‬‬

**Protein expression and purification:** Cells were grown at 37 °C in Luria-Bertani (LB) medium supplemented with ampicillin (100 µg/mL) for enzyme production. Protein expression was induced with 0.5 mM IPTG (G-Biosciences, St. Louis, USA) for five additional hours at 37 °C. Following induction, cells were harvested by centrifugation at 8000 × g for 10 minutes at 4 °C. The resulting cell pellets were resuspended in binding buffer (10 mM potassium phosphate buffer, 40 mM imidazole, and 500 mM NaCl, pH 7.4) and lysed by sonication under ice. Sonication was performed at 60 % amplitude in six cycles of 1-minute pulses, with 1-minute intervals between cycles. The cell lysate was centrifuged at 13,000 × g for 30 minutes at 4 °C to remove cell debris. The soluble fraction was separated and loaded onto a HisTrap™ HP column (GE Healthcare, Pittsburgh, USA) pre-equilibrated with binding buffer, following a previously described protocol^[1]^. Protein purity was confirmed by SDS-PAGE and western blot analysis using anti-His antibody (BioBharati, Kolkata, India). Protein concentrations were determined by measuring absorbance at 280 nm using a UV-vis spectrophotometer (Agilent Technologies, Santa Clara, USA). The modified Edelhoch and Gill/Von Hippel method available on ExPASy ProtParam was used for concentration calculations, assuming all cysteines were in a reduced state^[2]^. The calculated molecular weight and extinction coefficient of *Bs*EG2 (catalytic domain only) were 39565.17 kDa and 76,890 M^-1^ cm^-1^, respectively, while for *Bs*EG2 + CBM (catalytic domain + CBM), the values were 46928.24 kDa and 88350 M^-1^ cm^-1^, respectively.

**Determination of optimal temperature, pH, and stability:** The effects of temperature on *Bs*EG2 and *Bs*EG2 + CBM activity were evaluated using CMC as the substrate over a temperature range of 40 to 80 °C at a pH of 6.0. The optimum pH for both enzymes was determined using McIlvaine buffer across a pH range from 4 to 7, with *Bs*EG2 and *Bs*EG2 + CBM assayed at 55 °C and 60 °C, respectively. Thermal stability was assessed by measuring residual enzyme activity at 24-hour intervals at 55 °C, continuing until the specific activity dropped below 50% of the initial value. The half-life of *Bs*EG2 was calculated using the exponential decay equation, with enzyme activity prior to incubation set as 100 %. All enzyme assays were performed at their respective optimal temperatures and pHs.

**Enzyme activity assays:** *Bs*EG2 and *Bs*EG2 + CBM were assayed using 1 % sodium carboxymethyl cellulose (sourced from wood pulp, viscosity 1500-3000 cP, Na: 6.5-9.5 %) and β-glucan as substrates in McIlvaine buffer at pH_opt_ 6.0, with a total reaction volume of 150 µL at T_opt_ of 55 ℃. β-glucan from barley contains both β-1,3 and β-1,4 glycosidic linkages, which are found in natural cellulosic substrates, and was chosen as an alternative substrate due to its ability to be modulated between soluble and insoluble forms. Enzymatic hydrolysis was quantified using the DNS (3,5-dinitrosalicylic acid) assay, with glucose as the standard^[3]^. Following the completion of the enzymatic reaction, 150 µL of DNS reagent (1.3 M DNS, 1 M potassium sodium tartrate, and 0.4 N NaOH) was added to the reaction mixture and heated at 95 °C for 15 minutes in a dry bath. The assay mix was then centrifuged, and the absorbance of the supernatant was measured at 540 nm. All assays were conducted in triplicate and repeated at least three times, with standard deviations reported. One unit of enzymatic activity was defined as the amount of enzyme required to release one µmol of reducing sugar equivalents per minute. *Bs*EG2 specific activity was also assessed on additional substrates, including sugarcane bagasse, Avicel PH-101, filter paper, and lichenan.

**Kinetic analyses:** The kinetic parameters of *Bs*EG2 were determined using CMC and β-glucan as substrates. Enzymatic activity was measured under previously described reaction conditions. The reaction velocity was determined at various substrate concentrations ranging from 0.1-30 mg mL^-1^, using 0.03 µg of enzyme for a 30-minute reaction at T_opt_ (55 ℃) and pH_opt_ (6.0). Kinetic constants were calculated using a non-linear regression fit of the Michaelis-Menten equation in GraphPad PRISM version 8.0 (GraphPad Software, La Jolla, USA).

**Activity on biomass**: Enzyme activity was measured on sugarcane bagasse in its untreated form (SCB), and various pre-treated forms, including hemicellulose-free biomass (HFB), α-cellulose (AC), microcrystalline cellulose (MCC), and lignin-free biomass (LF). The preparations for these different forms have been previously reported^[4]^. To quantify the reducing sugars released, 1% (w/v) of each sugarcane bagasse type was used as the substrate and incubated with 5 µg of enzyme (*Bs*EG2 or *Bs*EG2 + CBM) in McIlvaine buffer (pH 6) at their respective optimal temperatures. The reactions were carried out under constant shaking for 2 hours. The amount of reducing sugars generated was determined using the DNS assay.

**Scanning electron microscopy (SEM):** Filter paper and sugarcane bagasse were hydrolyzed by *Bs*EG2 for 12 hours at 55 °C and subsequently analyzed using SEM. The reaction mixture had a final volume of 100 µL and contained 30 µg of enzyme in MES buffer (pH 6.0), along with 5 mg of bagasse or a disc of Whatman No. 1 filter paper as the substrate. Control reactions were prepared under identical conditions but without the enzyme. Following hydrolysis, the substrates were washed thrice with 500 µL Milli-Q water (Millipore, Merck, India), dried at 37 °C for 12 hours, and stored in a vacuum desiccator. The substrates were then coated with platinum using a Quorum 150R ES sputter coater (Quorum Technologies, Sussex, United Kingdom). Imaging was performed using a high-resolution environmental scanning electron microscope (Supra Max 55 V, Carl Zeiss AG, Jena, Germany), equipped with a field-emission gun (FESEM) Oxford 20 (Oxford, United Kingdom). Images were acquired under vacuum using a 5 kV accelerating voltage and a secondary electron detector. To ensure reproducibility, ten images were captured per sample from different areas.

**Effect of additives**: The impact of different ionic liquids, salts, and metal ions on the specific activity of *Bs*EG2 was evaluated by measuring enzyme activity in the presence of various concentrations of additives, as described in the enzyme activity assay above.

**Halotolerance and enzymatic activity in seawater:** The specific activity of *Bs*EG2 was measured in the presence of sodium chloride at concentrations up to 2.5 M. Seawater was collected from Puri, Orissa, India (19°47’44”N, 85°49’43”E), and filtered through a 0.22 µm membrane filter for further use. Enzymatic activity assays were conducted in two conditions: i) seawater with its pH adjusted to the enzyme’s optimal pH (pH 6.0) and ii) seawater at its natural pH (7.8) without any adjustment. The reactions were performed using 1% CMC as the substrate and 0.01% (v/v) enzyme at 60 °C. The reduced sugars were quantified using the DNS assay.

**Differential scanning fluorimetry:** The melting temperature (T_m_) of *Bs*EG2 in the presence of different additives was determined using Differential Scanning Fluorimetry (DSF)^[5]^. The 25 µL reaction mix contained 4.0 µg of enzyme in McIlvaine buffer, pH 6.0, 10X SYPRO™ Orange dye (Sigma-Aldrich, St. Louis, USA), and the respective additives. Fluorescence measurements were conducted in 8-well strips with optical caps (n = 4) over a 25–95 °C temperature range, with a ramp rate of 1 °C/min, using a real-time PCR system (Applied Biosystems). The apparent T_m_ was calculated by fitting the measured fluorescence data to a Boltzmann sigmoid equation, as described previously^[6]^.

**Cellulase processivity:** The processivity of *Bs*EG2 is the ratio of soluble reducing ends to insoluble reducing ends of sugars generated during filter paper assay reaction. Briefly, the reaction mixture consisted of 15 µg of BsEG2, 30 mg of Whatman filter paper (Grade 1), and McIlvaine buffer (pH 6.0). The mixture was incubated at 55 °C for 3 hours in a thermomixer (Eppendorf, Chennai, India). The reaction was terminated by heating at 95 °C for 10 minutes. After centrifugation at 13000g for 10 minutes, the supernatant was collected, and the reducing sugar content was quantified using the DNS assay with glucose as the standard. The insoluble pellet was washed three times with McIlvaine buffer (pH 6.0) to remove any residual soluble sugars^[7]^. The reducing ends of sugars in the insoluble fraction were then measured using the DNS assay.

**Multifunctionality assays:** Enzymatic assays were conducted using carboxymethyl cellulose (CMC), β-glucan from barley, *p*-nitrophenol (*p*NP)- based chromogenic substrates, including *p*-nitrophenol-β-D-cellobioside (*p*NPClb), *p*-nitrophenol-β-D-lactopyranoside (*p*NPLac), *p*-nitrophenol-β-D-maltoside (*p*NPMal), *p*-nitrophenol-β-D-galactopyranoside (*p*NPGal) and *p*-nitrophenol-β-D-glucopyranoside (*p*NPGlc). All assays were performed in McIlvaine buffer at pH_opt_ and incubated at *T*_opt_. Assays using *p*NP-based substrates following a previously reported protocol^[8]^. CMC and β-glucan hydrolysis products were quantified using the DNS assay, while glucose released from cellobiose hydrolysis was measured using the Glucose-oxidase-peroxidase assay (Glucose oxidase kit, Sigma-Aldrich, St. Louis, USA)^[9]^. To analyze the enzymatic hydrolysis products of cellooligosaccharides (cellobiose, cellotriose, cellotetraose, cellopentaose), reaction samples were collected at multiple time points and subjected to high-performance liquid chromatography (HPLC). The separation was performed using a Waters carbohydrate analysis column (3.9 × 300 mm, WAT084038) on a Waters HPLC system (Waters Corp., Milford, MA, USA) equipped with an evaporative light scattering (ELS) detector. The mobile phase consisted of a 3:1 ratio of acetonitrile and water. Oligosaccharide standards (glucose, cellobiose, cellotriose, cellotetraose, and cellopentaose) were used for peak identification and quantification.

For the complete conversion of oligosaccharides to glucose, the reaction products after 4 hours were quantified using the GOD-POD assay and further analyzed by thin-layer chromatography (TLC). TLC was performed using silica gel 60 F_254_ plates (Merck, Darmstadt, Germany). A 0.3 µL aliquot of the reaction product generated was spotted on the TLC plate, and the mobile phase consisted of n-butanol: glacial acetic acid: water (6:3:1). The spots were developed by spraying a freshly prepared mixture of aniline (0.2 g), diphenylamine (0.2 g) in acetone, and 2 mL of phosphoric acid (mixed immediately before use), followed by heating at 100 °C^[1]^.

**Molecular docking and molecular dynamics simulation:** We analyzed the evolutionary conservation of amino acid residues in the protein sequence of *Bs*EG2 by examining the phylogenetic relationships among homologous sequences using ConSurf server. Multiple sequence alignment (MSA) of these homologous sequences was performed using MAFFT. Based on predictions from AlphaFold-3, we generated the model structure of *Bs*EG2, focusing on residues 30 – 411. This model was created using the ColabFold implementation of AlphaFold, guided by the MSA. After refining the model structures of BsEG2 with and without the carbohydrate-binding module (CBM) through 10 ns of Molecular Dynamics (MD) simulations, we docked cellohexaose (G6) with these refined structures. The docking was conducted using the Deep Site and Docking Pose (DSDP) software platform, employing a blind docking approach. The docked structures were then sorted based on docking energy scores and root mean square deviation clustering. The G6 docked structures of *Bs*EG2, both with and without CBM, that exhibited the highest docking energy scores were selected for further MD simulation studies. All-atom MD simulations of the modelled structures of *Bs*EG2 and the G6 docked structures of *Bs*EG2 with and without CBM were performed using AMBER18^[10]^ . For MD simulation, the AMBER14SB and Glycam_06 force field parameters were considered for the protein and carbohydrate molecules, respectively ^[11]^. We analyzed average properties of the systems and binding interactions from the equilibrated trajectories using the CPPTRAJ module of AMBER18^[12]^.

Model systems were prepared using tLEAP, incorporating TIP3P water as an orthorhombic box with at least 15 Å padding around the solute. The systems were neutralized with randomly placed Na^+^ and Cl^−^ ions. Proteins and carbohydrates were parameterized with AMBER14SB and Glycam_06, respectively, while ions were parametrized using Joung & Cheatham parameters^[13]^. The systems were minimized for 20,000 steps (using steepest descent + conjugate gradient) under periodic boundary conditions. We heated the systems to 300 K over 100 ps NVT simulations with weak solute restraints using Langevin dynamics (friction coefficient 1 ps^−1^), followed by 300 ps NPT equilibration at 1 atm using the Berendsen barostat without restraints. Electrostatics were treated with Particle Mesh Ewald (PME)^[14]^, and covalent bonds to hydrogen were constrained using the SHAKE algorithm^[15]^ (tolerance 10^−8^ Å). Production MD simulations were conducted for 1 μs per system in the NPT ensemble using a 2 fs timestep and saving conformations every 1 ps for analysis.

**Inhibition studies using reaction products and substrates:** The enzymatic activity of *Bs*EG2 was assessed in the presence of its own hydrolysis products and cellobiose. Initially, a time-course enzymatic assay was performed by incubating *Bs*EG2 with CMC and β-glucan for specific time intervals. At each time point, the enzyme was heat-inactivated at 95 °C for 5 minutes, followed by the addition of an equal amount of fresh *Bs*EG2. The process was repeated at different time points to observe the impact of accumulated hydrolysis products on enzyme activity. In the next phase, reducing sugars generated from various substrates were added to fresh CMC, and the change in reaction rate upon addition of *BsEG2* was measured. Azo-CMC was used as a substrate to evaluate cellobiose inhibition, and increasing concentrations of exogenously added cellobiose (10 - 200 mM) were tested. The assay was performed under the optimum conditions of *Bs*EG2^[16]^.

**Synergistic interactions with other cellulases:** The synergy between *Bs*EG2 and β-glucosidase B8CYA8 (V169C/I246A/E173L) from *Halothermothrix orenii* was evaluated^[17]^. The assay was conducted in McIlvaine buffer, pH 6, at 60 ºC under a chosen set of assay conditions, ensuring that both enzymes exhibited 80-100% of their individual activities. The amount of glucose produced in the reaction was quantified using the Glucose Oxidase-Peroxidase (GOD-POD) assay (Glucose Oxidase Kit, Sigma-Aldrich, St. Louis, USA). To assess the synergistic action of *Bs*EG2 and B8CYA8 on Avicel to produce glucose, the degree of synergy was calculated using the following formula^[18]^:

$$Degree of synergy=\frac{Amount of product formed by the enzymes togetherly}{Amount of total product formed by the enzymes seperately}$$

**Figure S1.** **(a).** SDS-PAGE (12 %) image of purified *Bs*EG2. **(b)** SDS-PAGE (12 %) image of purified *Bs*EG2+CBM. Lane M is the PageRuler™ Plus Prestained Protein Ladder (Thermo Scientific, Waltham, USA). The proteins were stained by Coomassie Brilliant Blue R250.

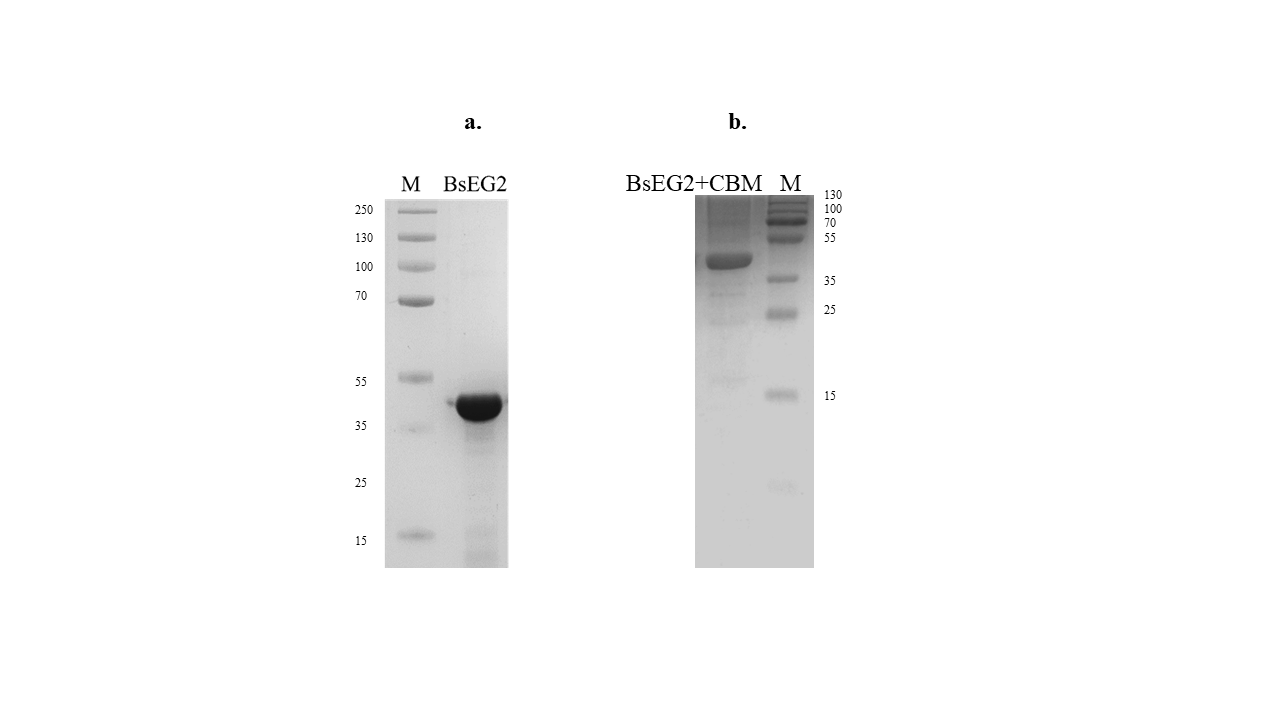

**Figure S2.** Multiple sequence alignment of *Bs*EG2 with other endoglucanases from the GH5 family. Conserved residues are highlighted, illustrating the sequence similarity and shared functional features within the enzyme family.

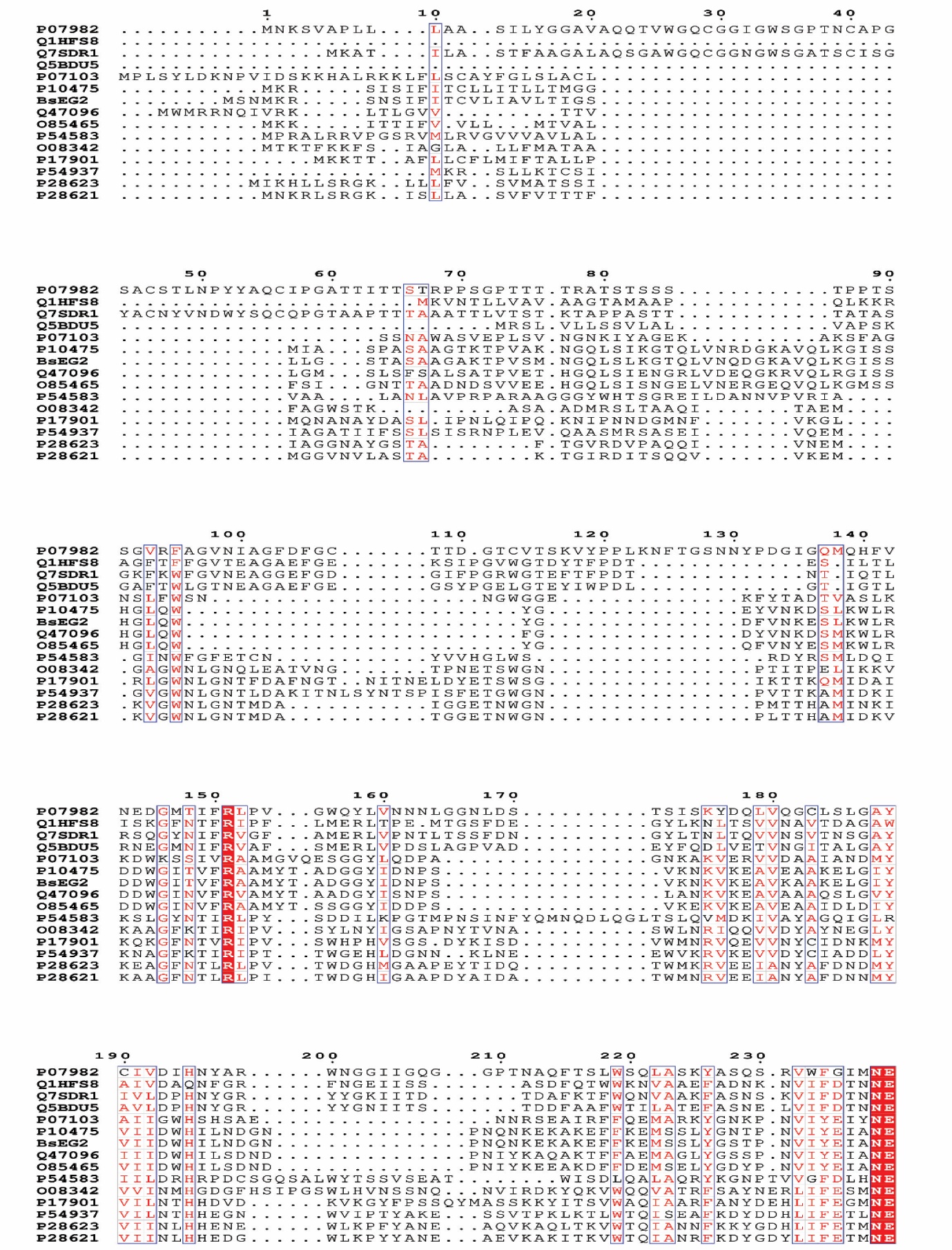

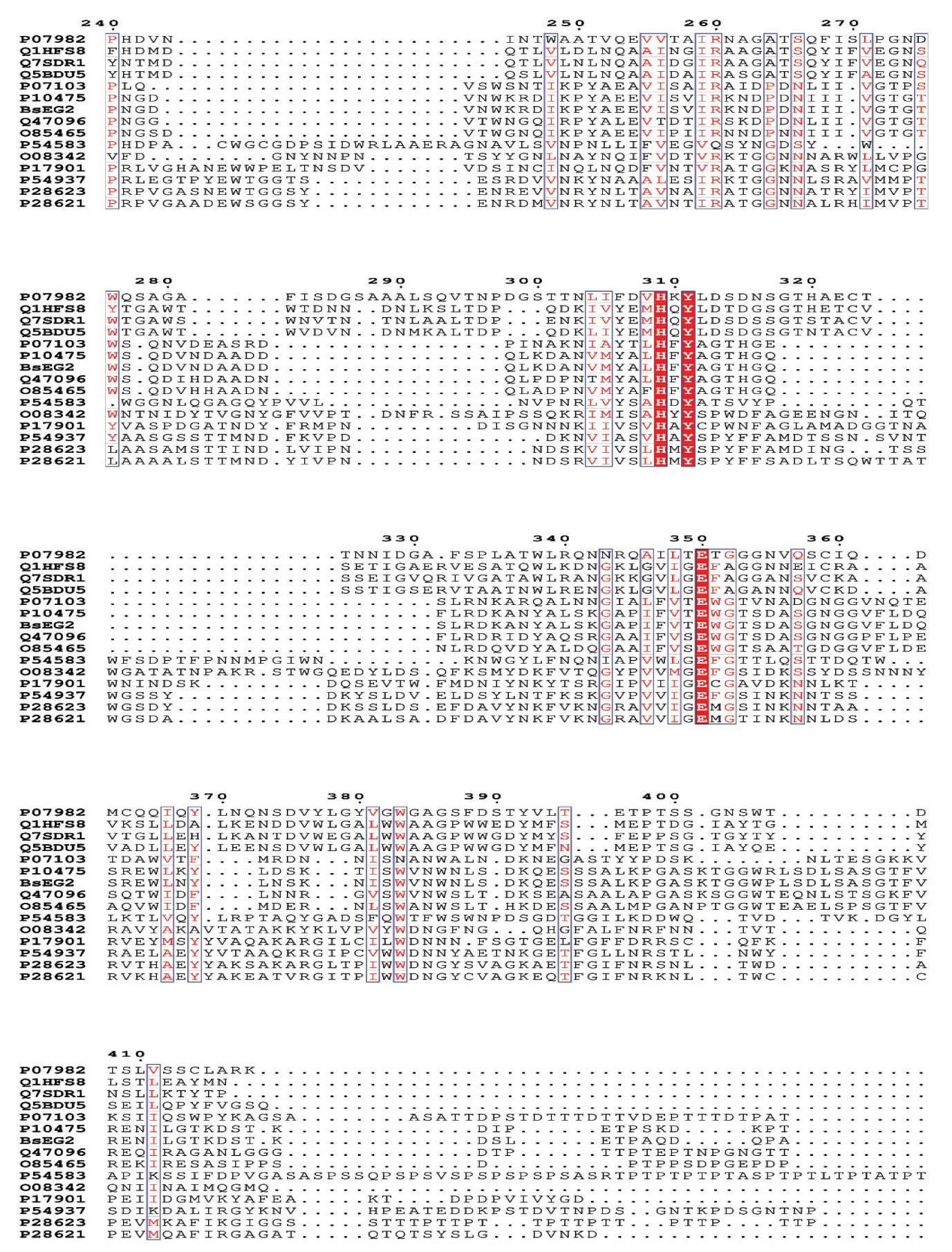

**Figure S3**. Heatmap of percentage identity matrix (PIM) for *Bs*EG2 and 14 other GH5 family endoglucanases. Colors represent sequence identity, ranging from low (blue) to high (red).

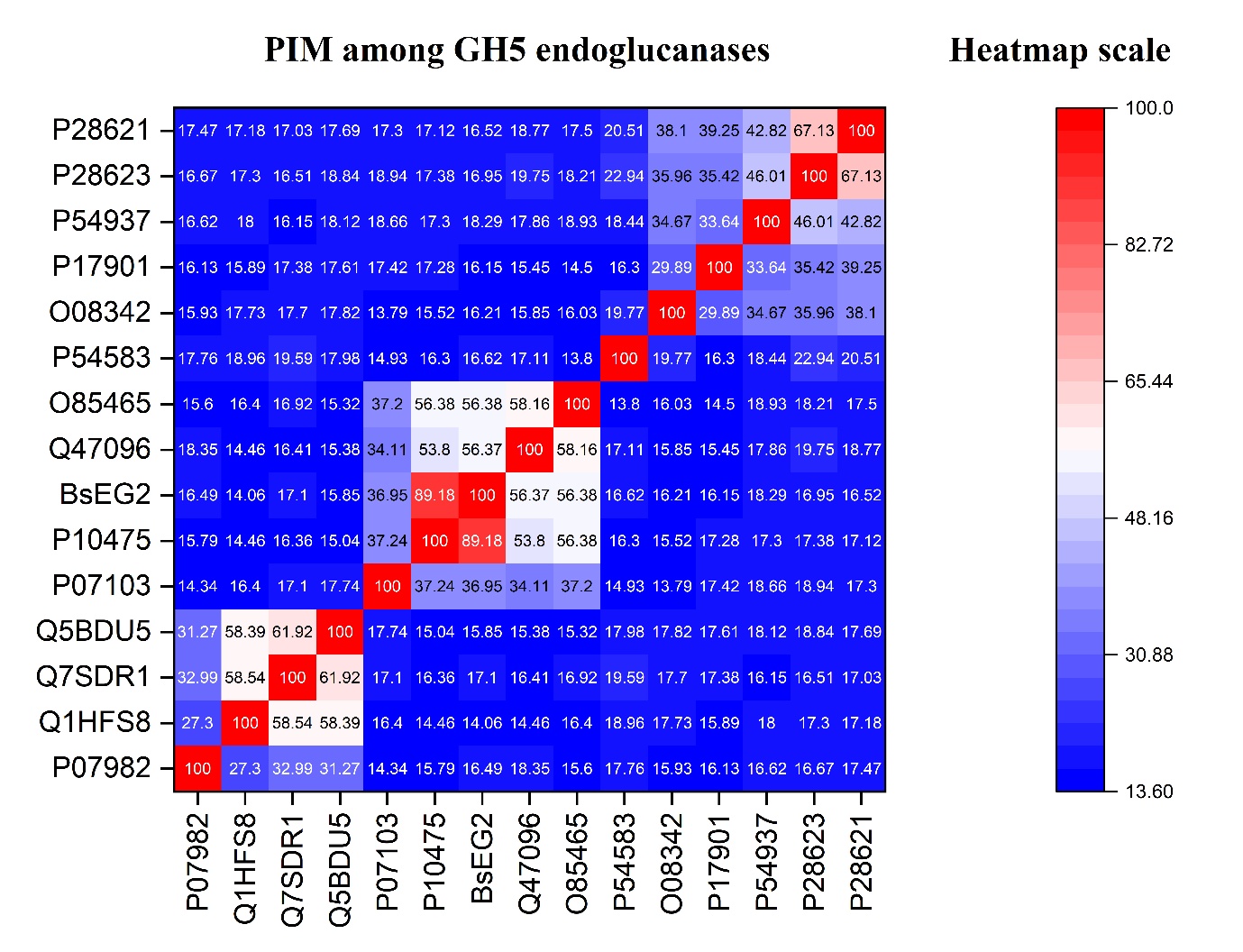

**Figure S4**. Determination of the optimal temperature of *Bs*EG2+CBM: The effect of temperature was measured over a range of 40 to 80 °C using 1% CMC as the substrate in McIlvaine buffer (pH 6.0). The concentration of reducing sugars generated was measured using the DNS assay and reported at different temperatures.

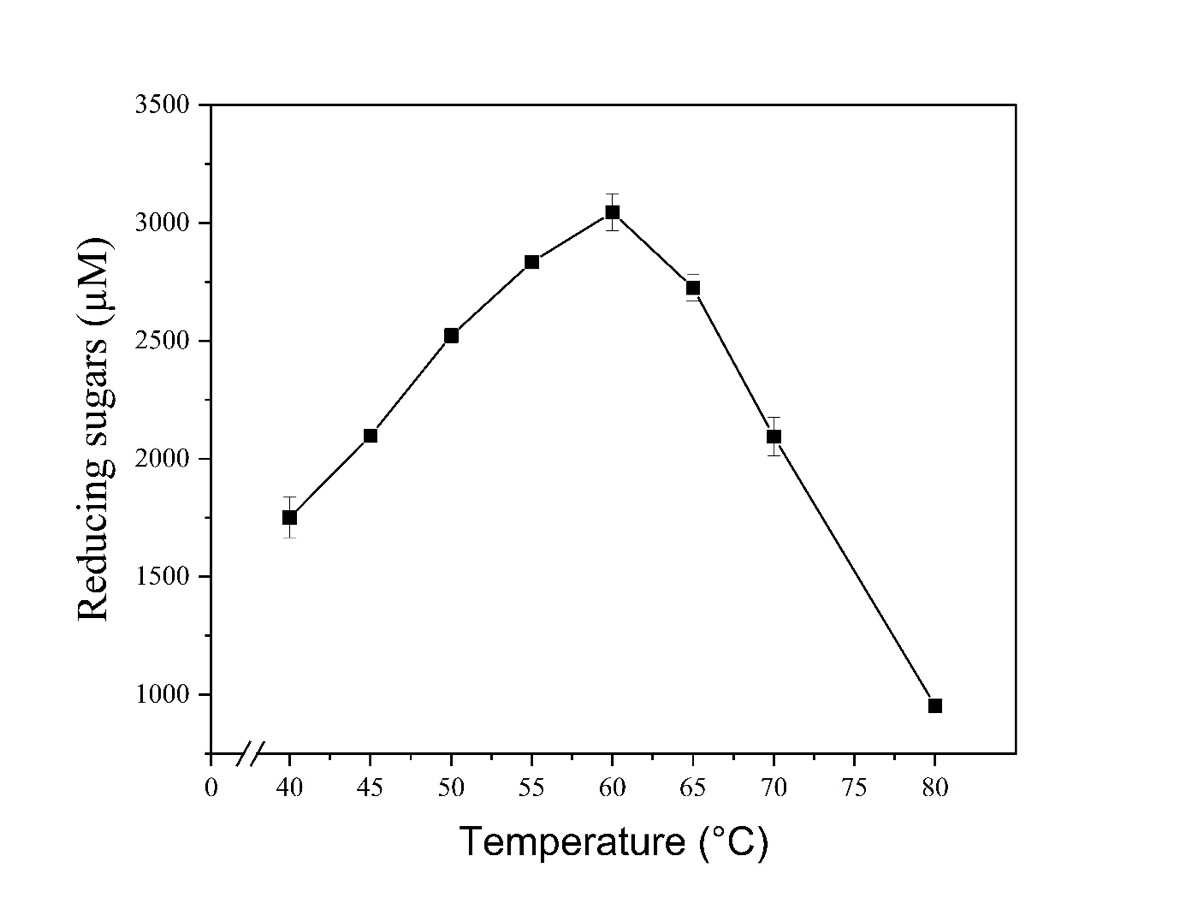

**Figure S5**. Determination of the optimal pH of *Bs*EG2+CBM: The effect of pH was measured over a range of pH 4 to pH 8, using 1% CMC as the substrate in McIlvaine buffer at 60 °C. The concentration of reducing sugars generated was measured by the DNS assay and reported against different pH levels.

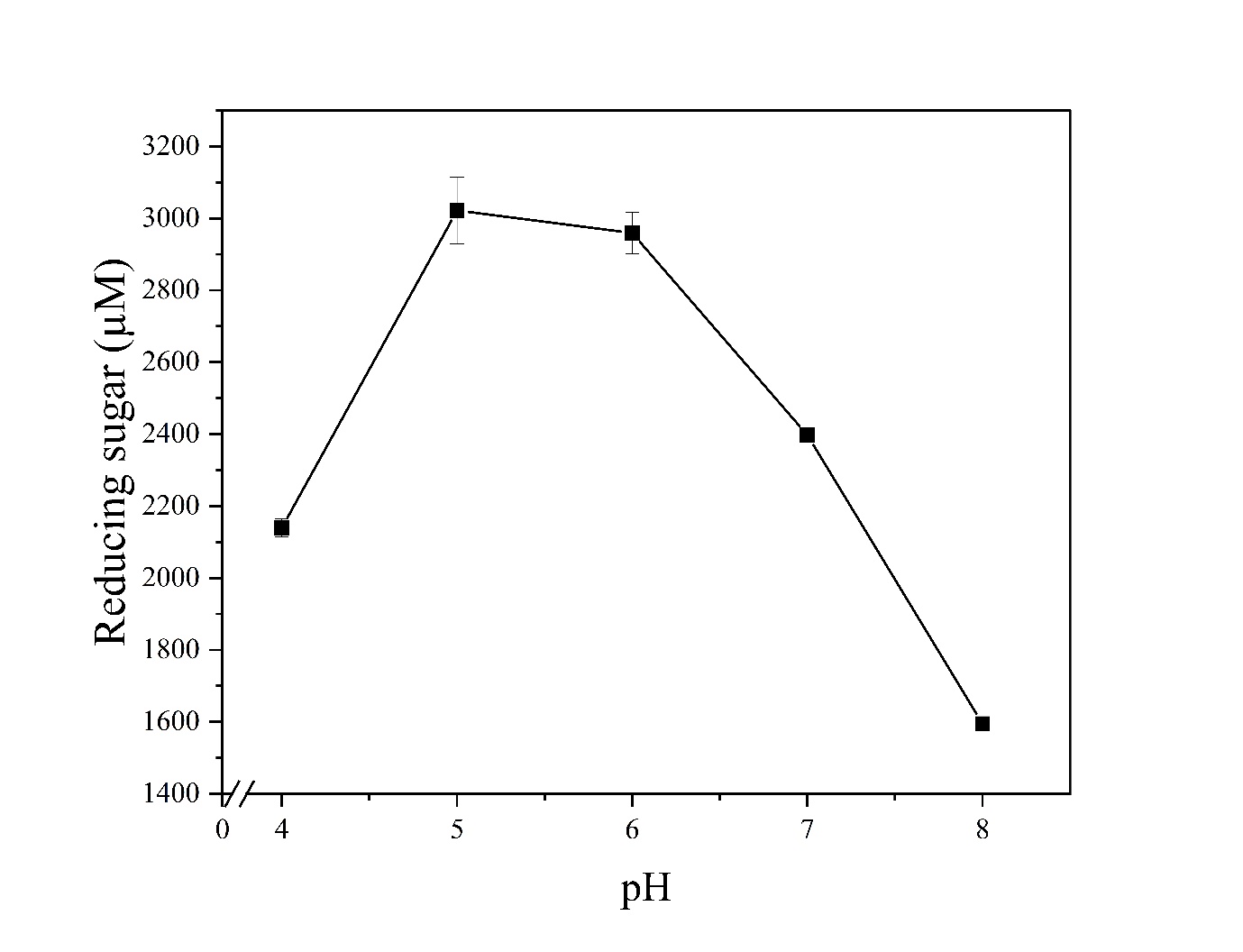

**Figure S6.** *Bs*EG2 catalyzed the hydrolysis of carboxymethyl cellulose. The *K_m_* and *V*_max_ were determined by a non-linear regression fit of the Michaelis–Menten equation using GraphPad Prism version 8.0. All experiments were done in triplicate and repeated at least three times. Errors shown are standard deviations of independent reactions.

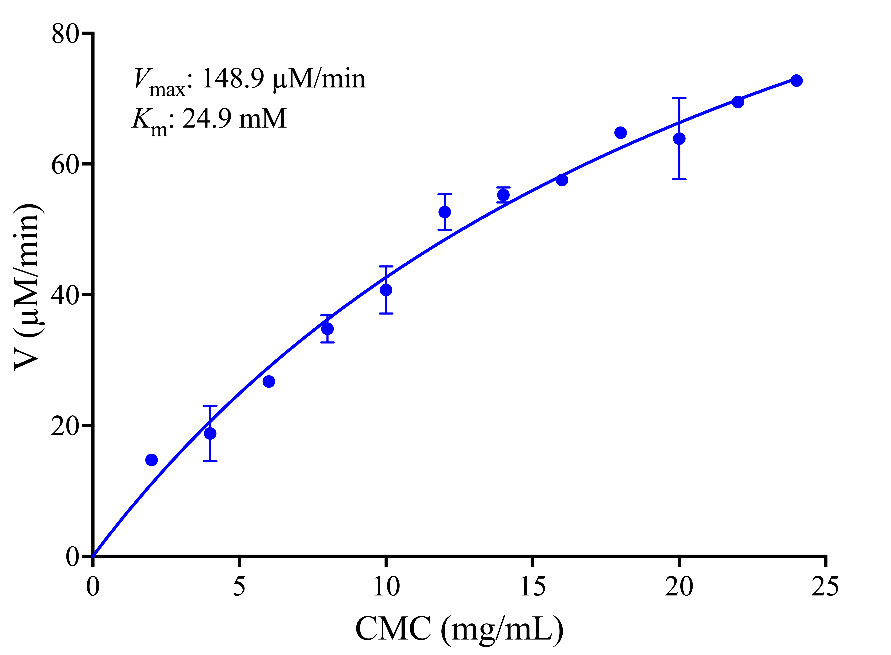

**Figure S7.** Scanning electron microscopy images showing the hydrolytic effect of *Bs*EG2 on filter paper and sugarcane bagasse. A - Scale bar of 10 µm for sugarcane bagasse and 3 µm for the filter paper, without enzyme. B - Scale bar of 1 µm for sugarcane bagasse and 300 nm for the filter paper, after incubation at 55 °C and MES buffer pH 6.0, in the presence of 30 μg *Bs*EG2 for 12 hours.

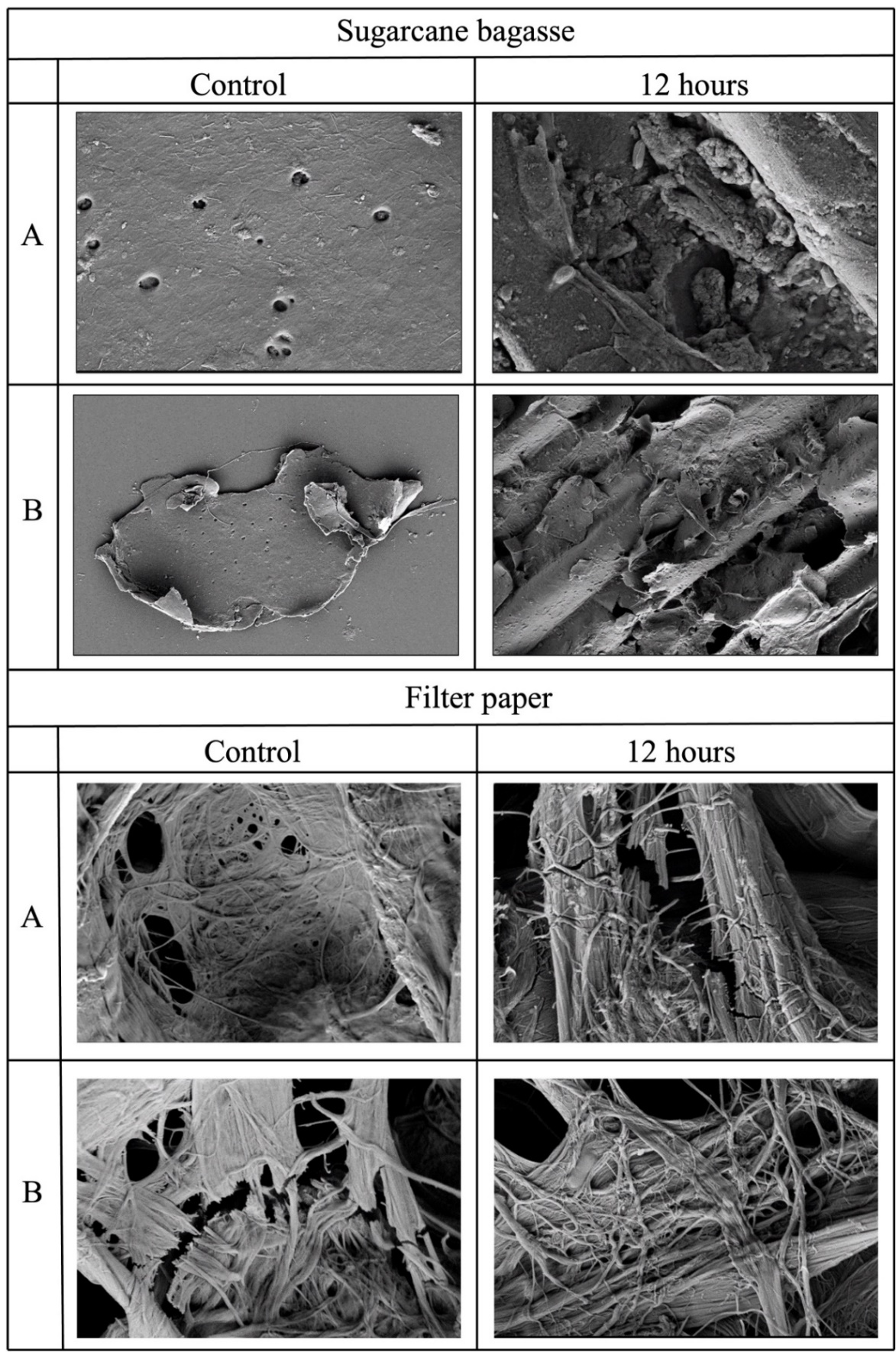

**Figure S8.** (a) AlphaFold predicted model structure of *Bs*EG2. (b)-(c) Simulated average G6 docked structure of *Bs*EG2 (green) superposed with initial docked structure (light pink) with and without CBM, respectively.

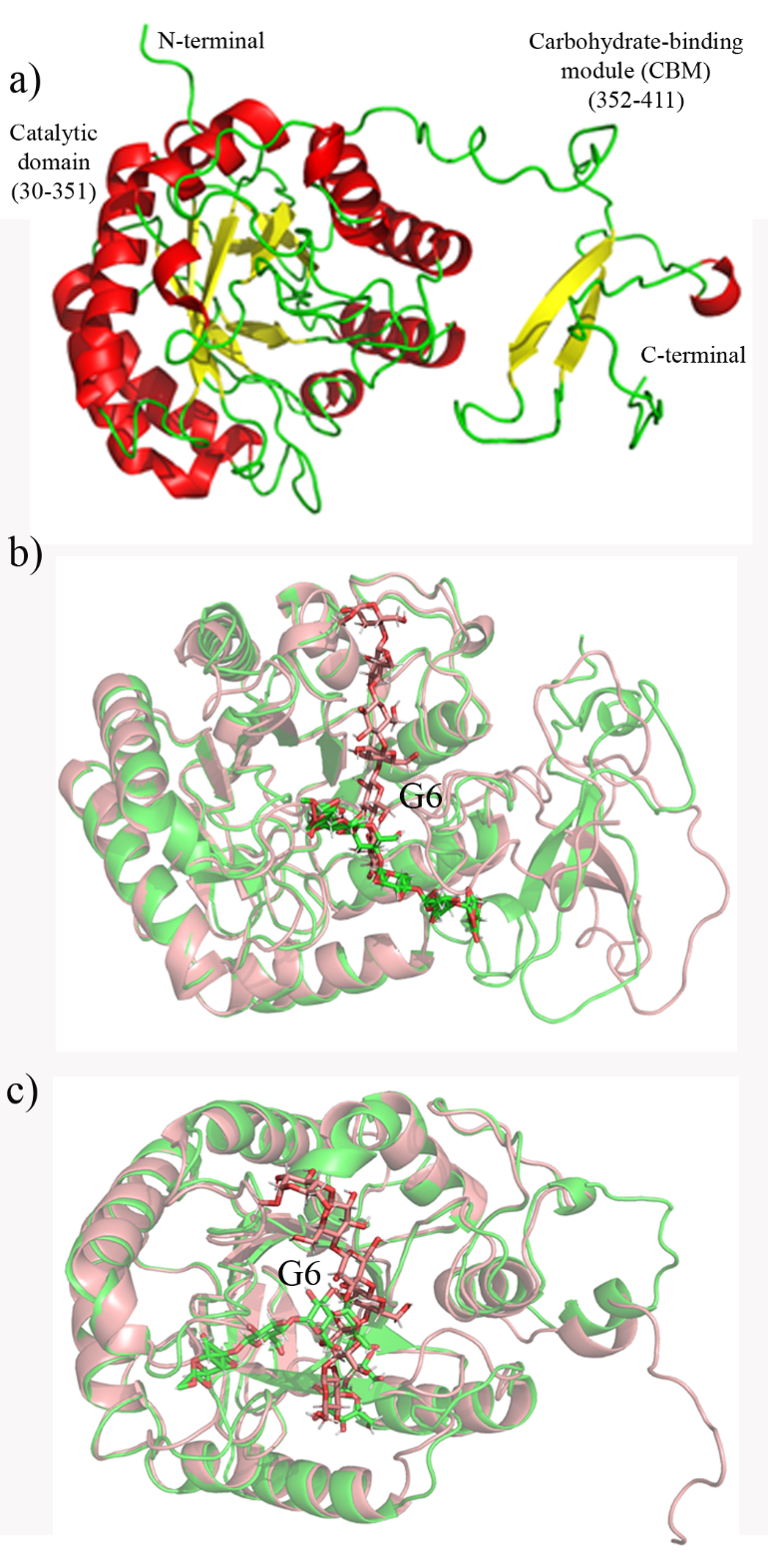

**Figure S9.** (a) RMSD of the *Bs*EG2 (green) and G6-bound BsEG2 (red) structure in the presence of CBM, and the *Bs*EG2 (yellow) and G6-bound BsEG2 (black) structure in the absence of CBM, with respect to their initial energy-minimized structures over the simulation time. (b) The mass center distance between the G6 and catalytic domain of *Bs*EG2 ($d_{G6-catalytic}$) in the absence (green) and presence (red) of CBM, and G6 and CBM of *Bs*EG2 ($d_{G6-CBM}$) in the presence of CBM (blue) over the simulation time. (c) In the *Bs*EG2-bound complex, the RMSF of the oligosaccharide moieties in G6 over the equilibrated MD trajectories *in the* presence (red) and absence (green) of CBM. (d) Distribution of the number of favorable polar hydrogen-bonded contacts between G6 and *Bs*EG2 ($N_{H}$) with (red) and without (blue) CBM.

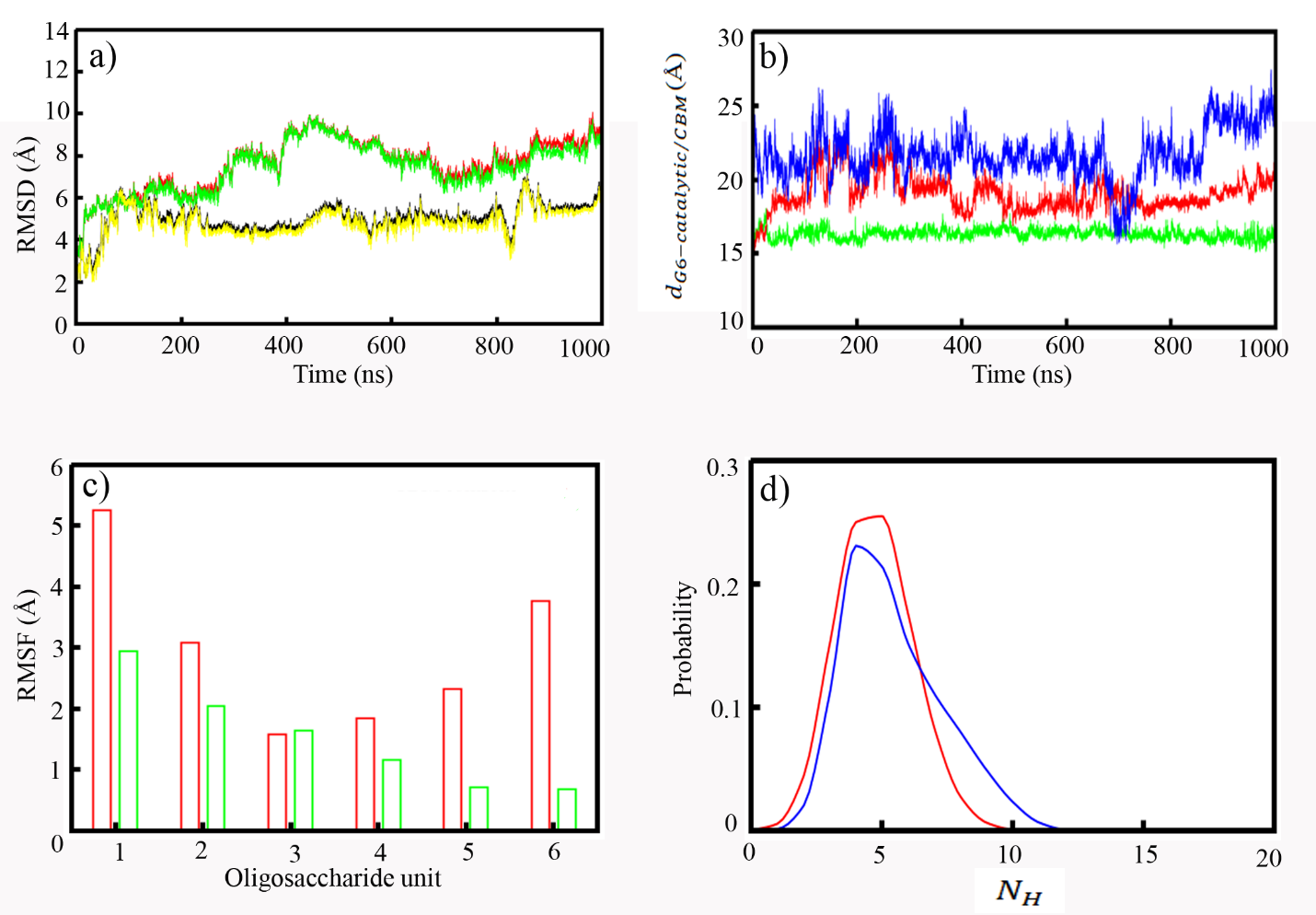

**Figure S10**. The evolutionary conservation of amino acid residues over the protein sequence of *Bs*EG2 based on the phylogenetic relations between homologous sequences.

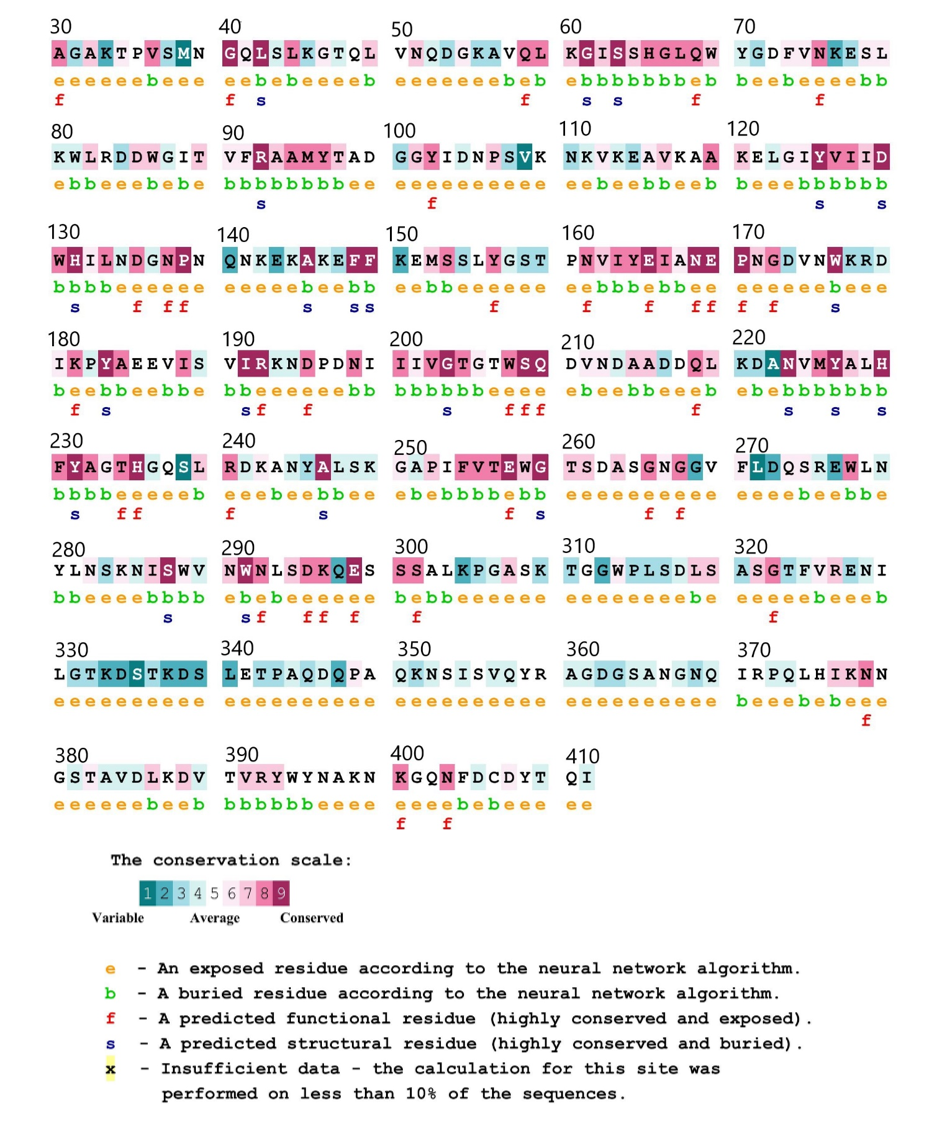

**Figure S11.** The impact of metal ions on *Bs*EG2. The % change in enzymatic activity in the presence of 10 mM metal ions for 30 minutes, at 55 °C and pH 6, is computed relative to the specific activity of *Bs*EG2 in the absence of metal ions, which is 340.57 ± 10.61 U/mg. The data is the average of three independent experiments run in triplicate. The error bars represent the standard deviation of independent experiments.

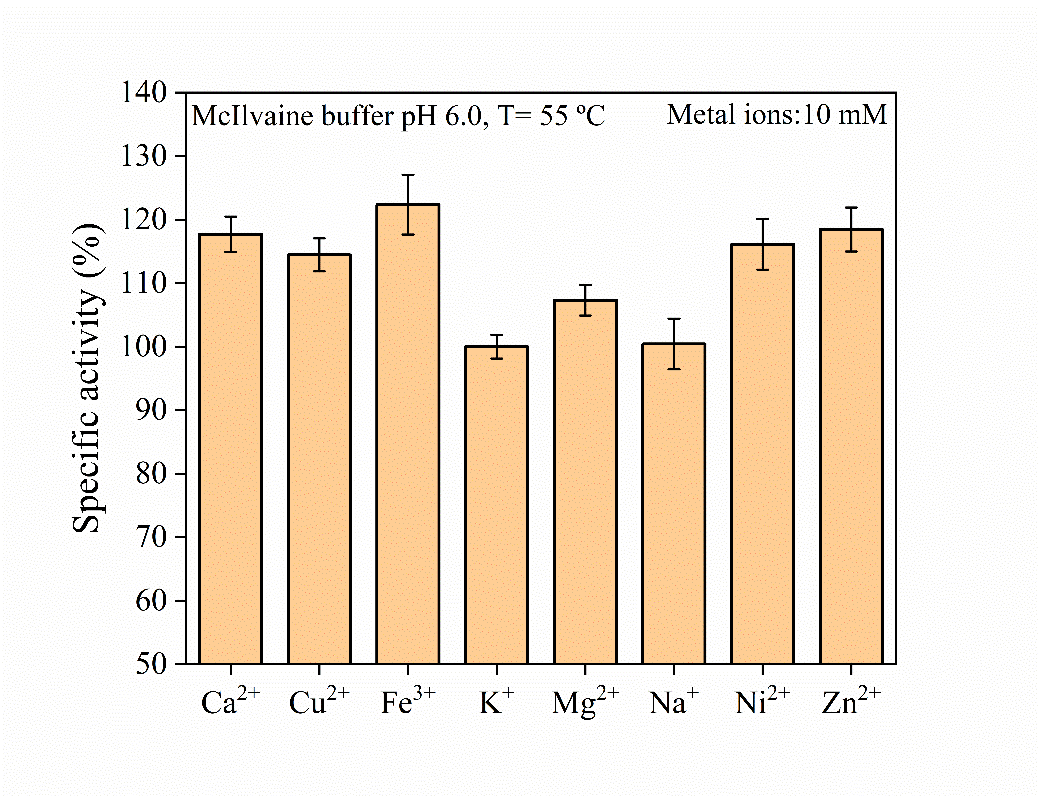

**Figure S12.** Hydrolysis product analysis of *Bs*EG2 using HPLC: The hydrolysis products of cellopentaose (G5) and cellotetraose (G4) by *Bs*EG2 were analyzed using high-performance liquid chromatography (HPLC) after 10 and 30 minutes of enzymatic reaction under optimal conditions. The enzymatic assay was terminated by heat inactivation at 95°C for 5 minutes. The hydrolysis products were identified based on retention times corresponding to glucose (G1), cellobiose (G2), cellotriose (G3), cellotetraose (G4), and cellopentaose (G5). The chromatographic analysis revealed that with an increase in reaction time, the concentration of accumulated cellobiose (G2) increased in both the cellotetraose and cellopentaose hydrolysis reactions. This indicates that cellobiose is the primary hydrolysis product, with some formation of cellotriose (G3).

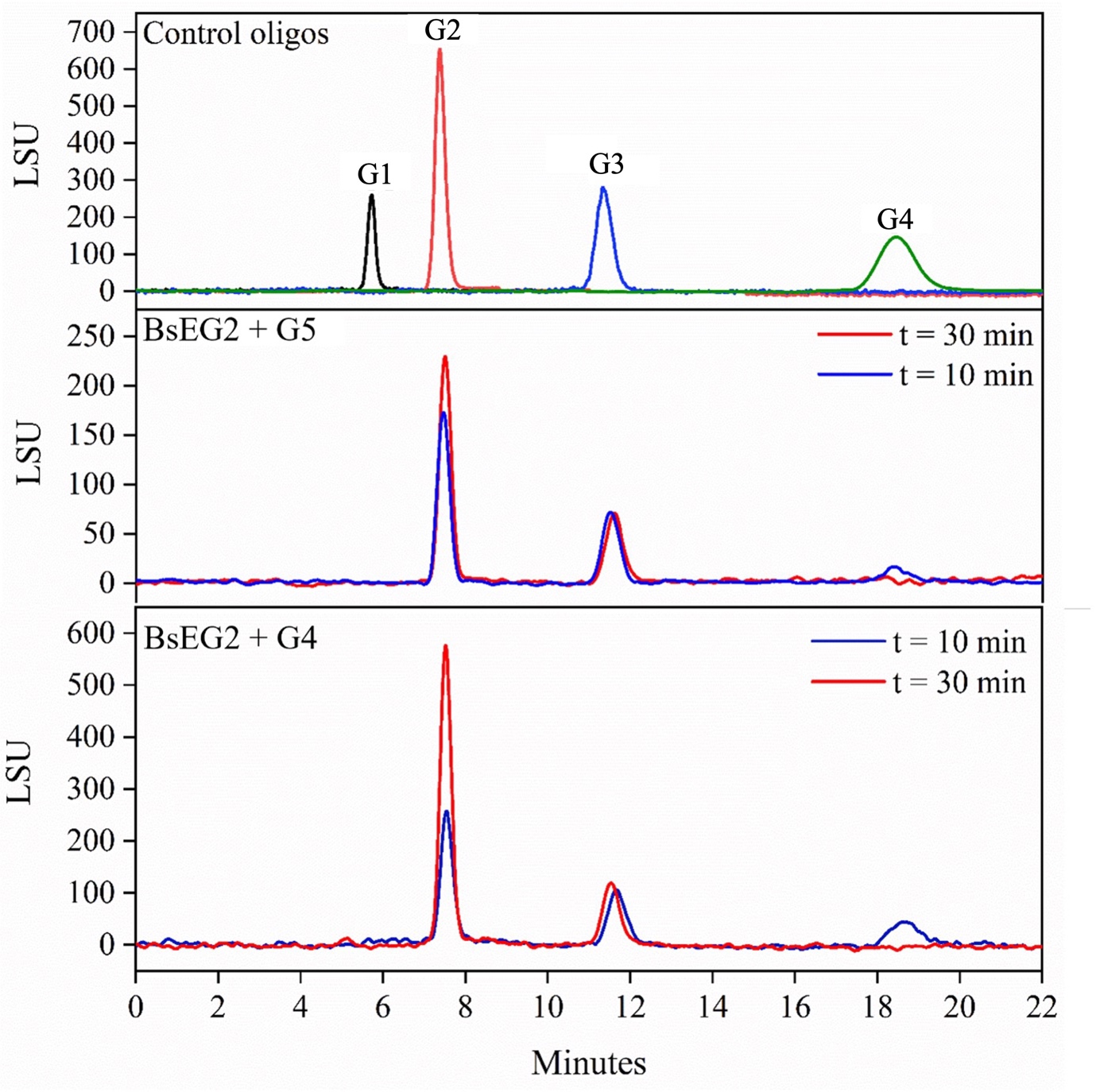

**Figure S13.** The identity of the final products of oligosaccharide hydrolysis by 10 μM BsEG2 was confirmed by TLC after a 4-h reaction at the optimum temperature and pH. The leftmost lane shows the reference spot for glucose (standard), followed by lanes L1 to L5, which display the hydrolysis products of oligosaccharides: cellobiose, cellotriose, cellotetraose, cellopentaose, and cellohexaose, respectively. The appearance of a single spot of glucose and the disappearance of spots for higher sugars indicate a 100% conversion of oligosaccharides to glucose.

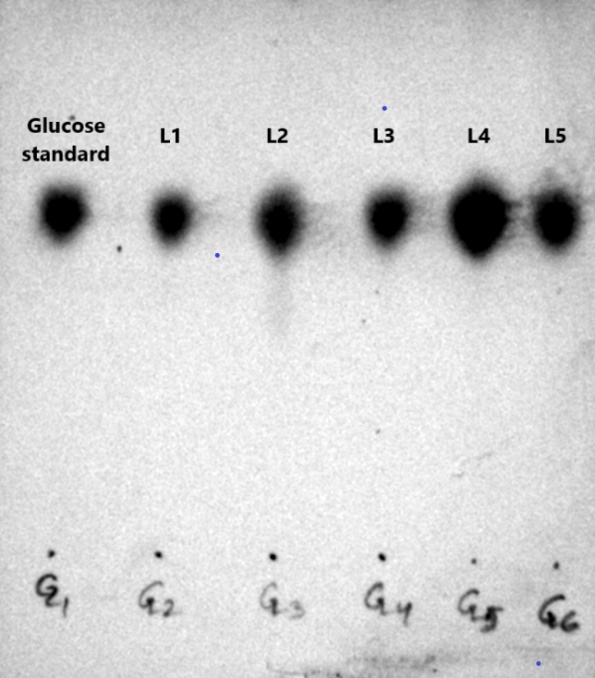

**Figure S14.** HPLC analysis of hydrolysis product of cellobiose by *Bs*EG2: Cellobiose was hydrolyzed by *Bs*EG2 under optimal conditions for 20 minutes. Following the enzymatic reaction, the enzyme was inactivated by heating the sample at 95°C for 5 minutes. The hydrolysis products were identified based on retention times corresponding to glucose and cellobiose. HPLC confirmed the conversion of cellobiose into glucose, demonstrating the enzymatic activity of *Bs*EG2 on cellobiose.

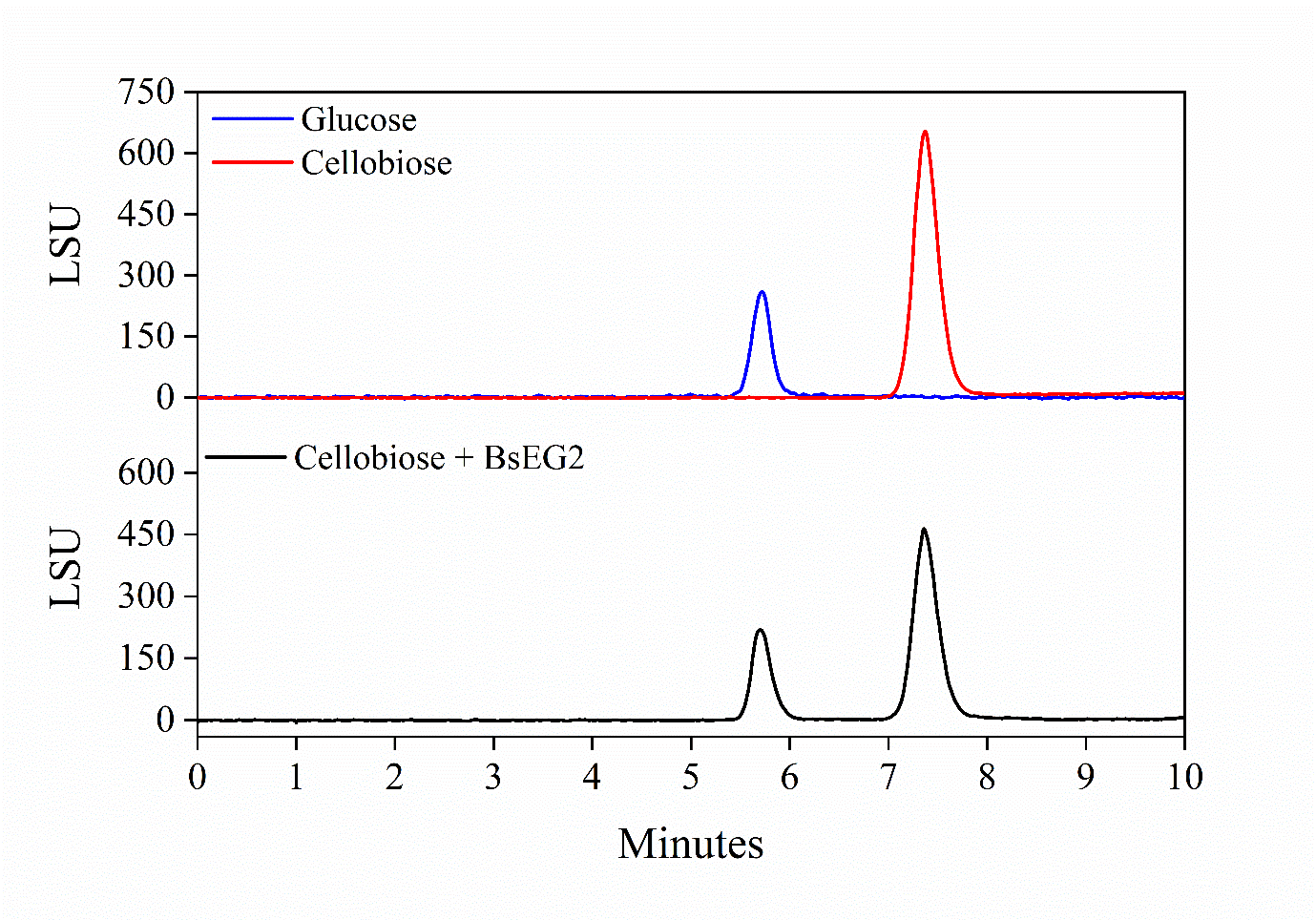

**Figure** **S15.** Specific activity was measured in the range of 10 to 200 mM cellobiose (Clb) using 0.1 µg BsEG2 and 1% azo-CMC as the substrate in McIlvaine buffer, pH 6.0, at 55 °C. The specific activity of *Bs*EG2 at T_opt_ and pH_opt_, in the absence of cellobiose, was set as 100%. Experiments were performed in triplicate and repeated at least three times.

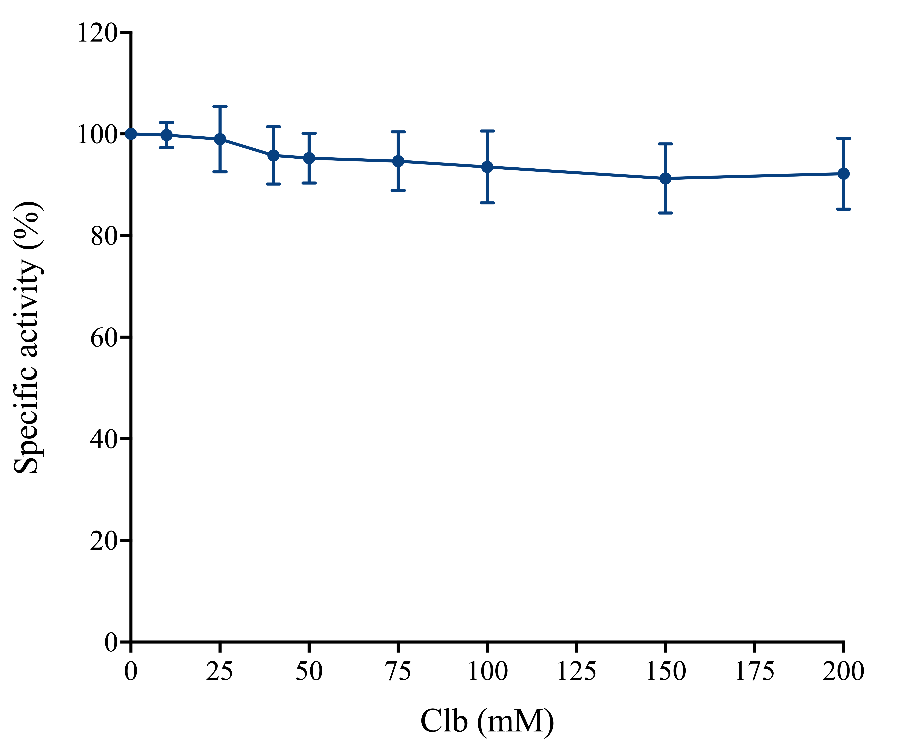

**Table S1.** Primers used in the cloning of *Bs*EG2 + CBM and *Bs*EG2

| **Primer name** | **Primer sequence** |
| --- | --- |
| *Bs*EG2 + CBM gene-specific forward primer | CGGAATTCATGATGCGAAGGAGGAAA |
| *Bs*EG2 + CBM gene-specific reverse primer | CGCTCGAGCTAATGGTGATGGTGA  TGGTGTCCAATCTGCGTGTAGTC |
| *Bs*EG2 gene-specific forward primer | CGGAATTCATGATGCGAAGGAGGAAA |
| *Bs*EG2 gene-specific reverse primer | CGCTCGAGCTAATGGTGATGGTGATGGTG  TGAACCACCACCACCTGAACCACCACCAC  CTGTGGGATTATCTTGTGC |

**Table S2**. Classification of the enzyme based on conserved sequences

| **Domain** | **Identifier sequence** | **Identifier ID** |
| --- | --- | --- |
| GH5 endoglucanase catalytic domain | GSTASAAGAKTPVSMNGQLSLKGTQLVNQDGKAV  QLKGISSHGLQWYGDFVNKDSLKWLRDDWGITVF  RAAMYTADGGYIDNPSVKNKVKEAVEAAKELGIY  VIIDWHILNDGNPNQNKEKAKEFFKEMSSLYGSTP  NVIYEIANEPNGDVNWKRDIKPYAEEVISVIRKND  PDNIIIVGTGTWSQDVNDAADDQLKDANVMYAL  HFYAGTHGQSLRDKANYALSKGAPIFVTEWGTS  DASGNGGVFLDQSREWLNYLNSKNISWVNWNL  SDKQESSSALKPGASKTGGWPLSD | PANTHER entry: PTHR34142 |
| CBM3 Carbohydrate binding domain | VQYRAGDGSANGNQIRPQLHIKNNGSTAVDLKDVTVRYWYNAKNKGQNFDCDYTQIGHHHHS | Prosite entry: PS51172 |

**Table S3.** Specific activity of *Bs*EG2 on various polysaccharides- β-glucan from barley, CMC, lichenan, Avicel-PH 101, sugar cane bagasse, filter paper, and chromogenic substrates- *p*NP-cellobioside, *p*NP-lactopyranoside, *p*NP-maltoside, *p*NP-galactopyranoside. Enzymatic activities were measured in McIlvaine buffer (pH 6.0) at 55 °C. Different amounts of purified BsEG2 were used depending on the substrate: 0.033 µg for CMC and β-glucan from barley, 0.32 µg for pNP-based chromogenic substrates, 0.64 µg for lichenan, 1.65 µg for Avicel-PH 101 and sugarcane bagasse-derived substrates, and 12 µg for filter paper. Incubation times were as follows:10 min for *p*NP-based substrates, 30 min for CMC and β-glucan from barley, 60 min for Avicel-PH 101 and natural substrates, and 2 hours for filter paper. Experiments were performed in triplicate and repeated at least three times. Error bars represent standard deviations from independent reactions. n.d. indicates that no detectable activity was observed.

| **Substrates** | **Specific Activity (µmoles min^-1^ mg^-1^)** |
| --- | --- |
| Artificial Substrates | |
| CMC-Na | 340.57 ± 10.61 |
| Avicel | 0.56 ± 0.04 |
| *p*NP-cellobioside | 62.50 ± 1.75 |
| *p*NP*-*lactopyranoside | 32.43 ± 1.03 |
| *p*NP-maltoside | n. d. |
| *p*NP-galactopyranoside | n. d. |
| Natural Substrates | |
| β-glucan (Barley) | 822.84 ± 16.73 |
| Lichenan | 10.45 ± 0.59 |
| Microcrystalline cellulose from bagasse | 14.70 ± 0.21 |
| Cellulose from bagasse | 0.77 ± 0.04 |
| Ground biomass from bagasse | 0.39 ± 0.02 |
| Hemicellulose-free biomass from bagasse | 0.61 ± 0.03 |
| Lignin-free biomass from bagasse | 1.09 ± 0.02 |
| Filter paper | 0.26 ± 0.01 |

n.d.: not detected

**Table S4**. Residues of *Bs*EG2 in the G6 binding site pocket as characterized by molecular docking and MD simulation studies, along with the energetics of G6 binding with *Bs*EG2 in the presence and absence of CBM.

| Interface residues of *Bs*EG2 in the G6 binding site | | | |
| --- | --- | --- | --- |
| Initial G6 docked structure with CBM | Initial G6 docked structure without CBM | Simulated average  G6 docked structure  with CBM | Simulated average G6 docked structure  without CBM |
| H65, G66, Q68, W69, Y70, Y96, D99, H131, L133, N168, E169, T206, W207, Q209, H229, F230, Y231, T234, H235, L239, E257, W258, G259, A263, S264, G265, W291, K296, E298, A365, N366 | H65, G66, Q68, W69, Y96, A98, D99, G100, G101, D104, H131, L133, N168, E169, T206, W207, Q209, H229, F230, Y231, T234, H235, E257, S261, A263, S264, G265, W291, N292, K296, E298, S299, S300 | **L133**, N134, D135, G136, N137, **E169**, N171, G172, D173,  W207, Q209, Y231,  **G233**, **T234**, H235, **D262**, A263, S264, Q350, S355, Q357, K377, N378, N379 | W69, R92, **Y96**, **D99**, H131, L133, **N134**, D135, G136, N168, E169, N171, T206, **W207**, S208, **Q209**, H229, F230, **Y231**, T234, L239, **E257**, A263, W291 |
| Energetics of G6 binding with *Bs*EG2 | | | |
| System | Interaction Energy (kcal/mol) | | Binding free energy (*ΔG*)  (kcal/mol) |
|  | Electrostatic  (*E_elec_*) | Van der Waals  (*E_vdw_)* |  |
| BsEG2 with CBM  + G6 | -88.82 ± 18.2 | -45.18 ± 6.4 | -13.15 ± 6.9 |
| BsEG2 without CBM + G6 | -90.67 ± 23.5 | -53.61 ± 4.9 | -21.56 ± 5.5 |
| Residues of *Bs*EG2 that make a favorable contribution to *ΔG* (kcal/mol) | | | |
| *Bs*EG2 with CBM + G6 | | *Bs*EG2 without CBM + G6 | |
| L133: -2.10 ± 0.7; N134: -1.14 ± 0.9; W207: -4.59 ± 0.9; T234: -2.36 ± 0.8; A263: -1.38 ± 0.7 | | L133: -1.59 ± 0.9; N134: -1.60 ± 0.9; W207: -2.81 ± 0.9; Q209: -1.86 ± 0.8; H229: -2.75 ± 0.8; F230: -3.10 ± 0.6; Y231: -4.0 ± 0.7; E257: -1.27 ± 1; A263: -1.10 ± 0.3; W291: -1.43 ± 0.6 | |

**Table S5.** Comparison of processivity (ratio of soluble sugars to insoluble sugars) generated from filter paper hydrolysis across various GH5 endoglucanases, including *Bs*EG2 from *Bacillus* sp. strain BS. The soluble sugars are designated as G1 (glucose), G2 (cellobiose), and G3 (cellotriose). The assay times are indicated where available.

| S. no. | Enzyme | Source | Processivity | Major products | Reference |
| --- | --- | --- | --- | --- | --- |
| 1. | CHU_2103 (3 h) | *Cytophaga hutchinsonii* | 3.72 | G2, G3 | ^[19]^ |
| 2. | HcCel5 | *Hahella chejuensis* KCTC 2396 | 5.75 | G2 | ^[20]^ |
| 3. | MtEG5A (1 h) | *Myceliophthora thermophile* | 3.91 | G2, G3 | ^[21]^ |
| 4. | EG5C (3 h) | *Bacillus subtilis BS-5* | 2.29 | G2, G3 | ^[22]^ |
| 5. | EG5C-1 (3 h) | *Bacillus subtilis BS-5* | 3.73 | G2, G3 | ^[22]^ |
| 6. | EG5C-2 (3 h) | *Bacillus subtilis BS-5* | 2.16 | G2, G3 | ^[22]^ |
| 7. | AS-HT-CeluzA (3 h) | *A. ochraceus* MTCC1810 | 1.79 | N.D. | ^[23]^ |
| 8. | Cel5H | *S. degradans* | 4.25 | G2 | ^[24]^ |
| 9. | *Bs*EG2 (3 h) | *Bacillus* sp. strain BS | 2.65 | G1, G2 | This study |

N.D. : Not determined

**References**

[1]	S. Goswami, N. Gupta, S. Datta, *Biotechnology for Biofuels* **2016**, *9*, 72.

[2]	aE. Gasteiger, C. Hoogland, A. Gattiker, S. e. Duvaud, M. R. Wilkins, R. D. Appel, A. Bairoch, in *The Proteomics Protocols Handbook* (Ed.: J. M. Walker), Humana Press, Totowa, NJ, **2005**, pp. 571-607; bS. Duvaud, C. Gabella, F. Lisacek, H. Stockinger, V. Ioannidis, C. Durinx, *Nucleic Acids Research* **2021**, *49*, W216-W227.

[3]	G. L. Miller, *Analytical Chemistry* **1959**, *31*, 426-428.

[4]	A. Konar, S. Aich, R. Katakojwala, S. Datta, S. V. Mohan, *Appl Microbiol Biotechnol* **2022**, *106*, 6059-6075.

[5]	F. H. Niesen, H. Berglund, M. Vedadi, *Nature Protocols* **2007**, *2*, 2212-2221.

[6]	S. K. Sinha, S. Datta, *Applied Microbiology and Biotechnology* **2016**, *100*, 8399-8409.

[7]	R. Pan, Y. Hu, L. Long, J. Wang, S. Ding, *Enzyme and Microbial Technology* **2016**, *91*, 42-51.

[8]	S. Aich, R. K. Singh, P. Kundu, S. P. Pandey, S. Datta, *Biotechnol Biofuels* **2017**, *10*, 135.

[9]	S. K. Sinha, M. Datta, S. Datta, *Green Chemistry* **2021**, *23*, 7299-7311.

[10]	I. Y. B.-S. D. A.Case, S. R.Brozell, D. S.Cerutti, T. E.Cheatham, III, V. W. D.Cruzeiro, T. A.Darden, R. E.Duke, D.Ghoreishi, M. K.Gilson, H.Gohlke, A. W.Goetz, D.Greene, R.Harris, N.Homeyer, Y.Huang, S.Izadi, A.Kovalenko, T.Kurtzman, T. S.Lee, S.LeGrand, P.Li, C.Lin, J.Liu, T.Luchko, R.Luo, D. J.Mermelstein, K. M.Merz, Y.Miao, G.Monard, C.Nguyen, H.Nguyen, I.Omelyan, A.Onufriev, F.Pan, R.Qi, D. R.Roc, A.Roitberg, C.Sagui, S.Schott-Verdugo, J.Shen, C. L.Simmerling, J.Smith, R.Salomon-Ferrer, J.Swails, R. C.Walker, J.Wang, H.Wei, R. M.Wolf, X.Wu, L.Xiao, D. M.York, and P. A.Kollman, AMBER2018, University of California, San Francisco, CA, 2018., University of California, San Francisco, CA, **2018**.

[11]	aV. Hornak, R. Abel, A. Okur, B. Strockbine, A. Roitberg, C. Simmerling, *Proteins* **2006**, *65*, 712-725; bK. N. Kirschner, A. B. Yongye, S. M. Tschampel, J. González-Outeiriño, C. R. Daniels, B. L. Foley, R. J. Woods, *Journal of computational chemistry* **2008**, *29*, 622-655.

[12]	D. R. Roe, T. E. Cheatham, 3rd, *Journal of chemical theory and computation* **2013**, *9*, 3084-3095.

[13]	I. S. Joung, T. E. Cheatham, 3rd, *The journal of physical chemistry. B* **2008**, *112*, 9020-9041.

[14]	T. Darden, D. York, L. Pedersen, *The Journal of Chemical Physics* **1993**, *98*, 10089-10092.

[15]	R. Elber, A. P. Ruymgaart, B. Hess, *The European physical journal. Special topics* **2011**, *200*, 211-223.

[16]	M. Gruno, P. Väljamäe, G. Pettersson, G. Johansson, *Biotechnology and Bioengineering* **2004**, *86*, 503-511.

[17]	S. K. Sinha, S. Das, S. Konar, P. K. Ghorai, R. Das, S. Datta, *International Journal of Biological Macromolecules* **2020**, *156*, 621-632.

[18]	J. Woodward, *Bioresource Technology* **1991**, *36*, 67-75.

[19]	C. Zhang, Y. Wang, Z. Li, X. Zhou, W. Zhang, Y. Zhao, X. Lu, *Applied Microbiology and Biotechnology* **2014**, *98*, 6679-6687.

[20]	S. Wu, S. Wu, *Applied Biochemistry and Biotechnology* **2020**, *190*, 448-463.

[21]	A. Karnaouri, M. N. Muraleedharan, M. Dimarogona, E. Topakas, U. Rova, M. Sandgren, P. Christakopoulos, *Biotechnol Biofuels* **2017**, *10*, 126.

[22]	B. Wu, S. Zheng, M. M. Pedroso, L. W. Guddat, S. Chang, B. He, G. Schenk, *Biotechnology for Biofuels* **2018**, *11*, 20.

[23]	P. Asha, J. Divya, I. S. Bright Singh, *Bioresour Technol* **2016**, *213*, 245-248.

[24]	S. S. Ghatge, A. A. Telke, S.-H. Kang, V. Arulalapperumal, K.-W. Lee, S. P. Govindwar, Y. Um, D.-B. Oh, H.-D. Shin, S.-W. Kim, *Applied Microbiology and Biotechnology* **2014**, *98*, 4421-4435.
